# Experimental assay of a fitness landscape on a macroevolutionary scale

**DOI:** 10.1101/222778

**Authors:** Victoria O. Pokusaeva, Dinara R. Usmanova, Ekaterina V. Putintseva, Lorena Espinar, Karen S. Sarkisyan, Alexander S. Mishin, Natalya S. Bogatyreva, Dmitry N. Ivankov, Arseniy V. Akopyan, Sergey Ya. Avvakumov, Inna S. Povolotskaya, Guillaume J. Filion, Lucas B. Carey, Fyodor A. Kondrashov

**Author notes:** Equal contribution (Victoria O. Pokusaeva, Dinara R. Usmanova, Ekaterina V. Putintseva).

## Abstract

Characterizing the fitness landscape, a representation of fitness for a large set of genotypes, is key to understanding how genetic information is interpreted to create functional organisms. Here we determined the evolutionarily-relevant segment of the fitness landscape of His3, a gene coding for an enzyme in the histidine synthesis pathway, focusing on combinations of amino acid states found at orthologous sites of extant species. Just 15% of amino acids found in yeast His3 orthologues were always neutral while the impact on fitness of the remaining 85% depended on the genetic background. Furthermore, at 67% of sites, substitutions are under sign epistasis, having both strongly positive and negative effect in different genetic backgrounds. 46% of sites were under reciprocal sign epistasis. Sign epistasis affected few genotypes but involved interaction of multiple sites, shaping a rugged fitness landscape in which many of the shortest paths between highly fit genotypes are inaccessible.

---

Predicting function and fitness of organisms from their genotypes is the ultimate goal of many fields in biology, from medical genetics to systems biology to the study of evolution^1–5^. Among the conceptual frameworks for understanding the genotype to phenotype connection is the fitness landscape, which assigns a fitness (phenotype) to every possible genotype (sequence) of a gene or genome under consideration^4,6^. The recognition of the importance of the fitness landscape stimulated the development of a variety of theoretical approaches to its description, including its general shape and epistatic interactions between alleles, a key property which determines the complexity of the fitness landscape (see [ref. 4] and references within). Before the advent of next-generation sequencing, experimental assays of the fitness landscape were few and could not address the issue at the sequence level. Recently, large-scale experimental assays described the shape of the fitness landscape a few mutations away from a local fitness peak (see [ref. 7-10] and references within). Also, some assays involving a smaller number of genotypes considered combinations of mutations with established functional ^11–16^ or evolutionary ^17–22^ significance.

Empirical evidence of the nature of large-scale fitness landscapes mostly comes from the study of genotypes incorporating random mutations^4,7–10^, the majority of which are deleterious^7- 10,23^. Thus, our present knowledge of fitness landscapes is primarily driven by the study of deleterious mutations and their interactions, although local adaptive trajectories have also been considered^2,4,16,24–26^. Deleterious mutations were found to engage in synergistic epistasis, whereby the joint effect of multiple mutations was stronger than the sum of their individual effects^4,7–10,16^. Furthermore, sign epistasis among random mutations was mostly rare^5,7–10,16,27^, although some of these conclusions differ from study to study *(e.g.* see [4,27]).

Unfortunately, there are fundamental limitations to assaying the fitness landscape on a large or macroevolutionary level with random mutation libraries. The number of genotypes underlying the fitness landscape is the combinatorial set of all amino acids across the length of the protein^4,6^. For example, for the 220 amino acid protein coded by the His3 gene in *Saccharomyces cerevisiae,* the fitness landscape is a 220 dimensional genotype space with 20^220^ different possible sequences. Such immense spaces are both computationally and experimentally intractable. Fortunately, it may not be necessary to survey all genotypes to study the evolutionary-relevant section of the fitness landscape. Because the vast majority of mutations in protein sequences are deleterious^23^, a randomly sampled protein sequence is non-functional^28,29^.

Here we propose an evolutionary approach for assaying fitness landscapes on a macroevolutionary scale in a high-throughput manner that avoids the random sampling of mostly non-functional sequences. The functionally and evolutionarily relevant section of the fitness landscape can be represented by the combination of extant amino acid states, those found in extant species. This approach, applied previously on a limited scale^17–22^ mitigates the problem of exploring a prohibitively large fitness landscape while highlighting the relationships between evolutionarily-relevant genotypes **(Fig. 1a)**. Crucially, substitutions that have been fixed in evolution are fundamentally different from random mutations, the former are either neutral or beneficial in at least some genetic contexts and represent the driving force of molecular evolution, while the latter are mostly deleterious and are primarily relevant on a microevolutionary scale. Therefore, current empirical data do not shed much light on the impact of interactions between substitutions that fixed in the course of evolution by natural selection. Combinations of extant amino acid states allow one to assay a much wider functionally relevant area of the sequence space than approaches based on random mutagenesis of a single sequence (**Fig. 1b,c**).

**Figure 1.**
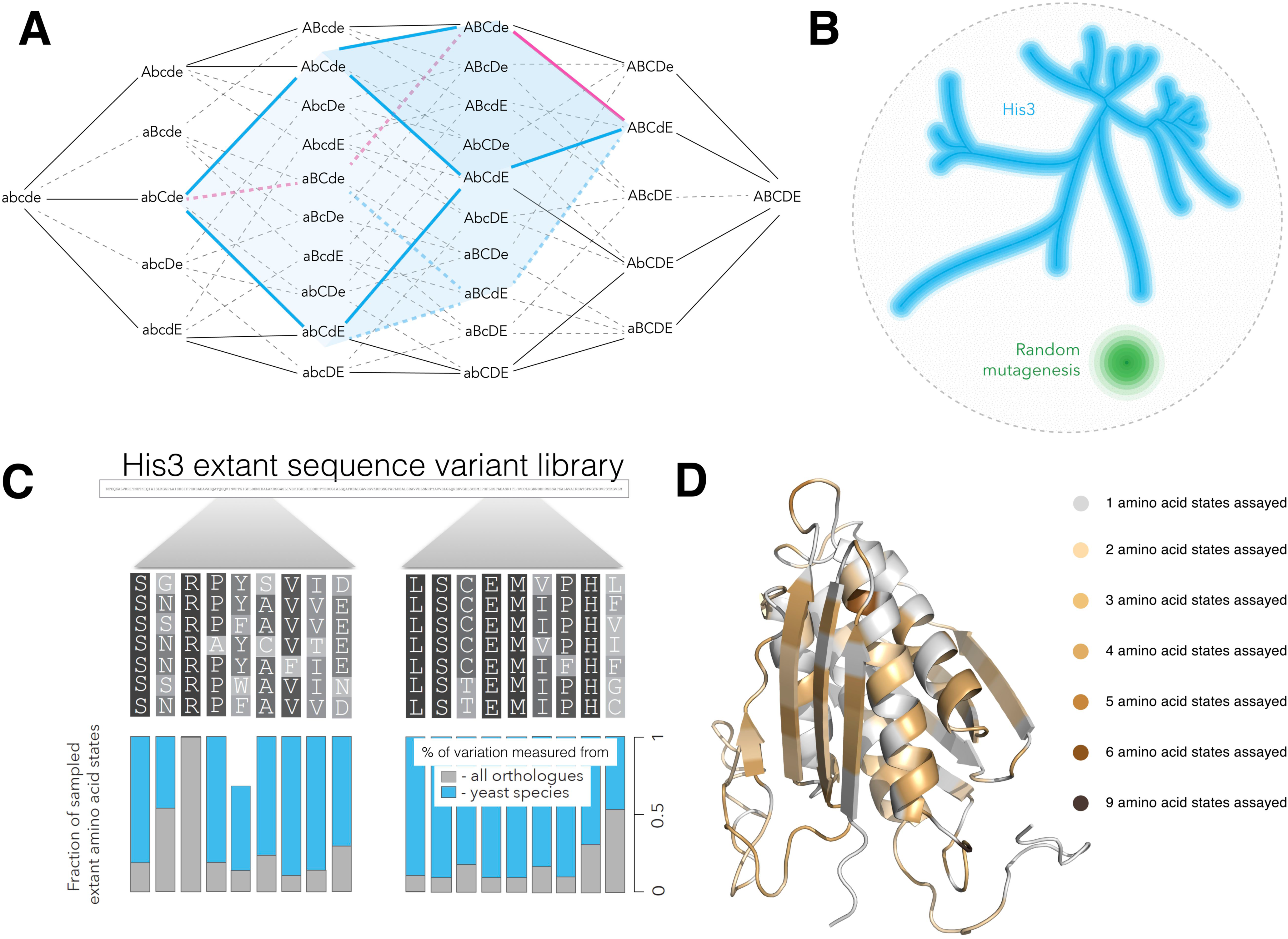
Combinatorial approach to the study of fitness landscapes. A fitness landscape is the representation of fitness for all possible genotypes composed of a specific set of loci. **a,** Following Figure 1 from Sewall Wright ref. [6] consider the genotype space consisting of 5 loci, each with two allele states (lower and uppercase letters). The entire genotype space is 5-dimensional consisting of 2^5^ genotypes. Given two genotypes found in extant species (abCde and ABCdE in this example), surveying combinations of extant alleles substantially reduces the dimensionality of the genotype space, concomitantly reducing the number of genotypes to assay. The surveyed area (blue cube) considers all combinations of allele substitutions that have occurred in the course of evolution between the two sequences (red line), avoiding the sampling of combinations with less relevance to the evolutionary trajectory (black lines). **b,** Given the entire multidimensional genotype space (black circle) our approach considers an multidimensional subspace consisting of the combinatorial set of amino acid states from extant species. The blue line represents the yeast phylogeny and the surrounding blue space represents a multidimensional set of combinations of extant amino acids of the sequence under consideration, one His3 gene segment in our study. By contrast, random mutagenesis studies consider only a local segment of the genotype space surrounding a specific genotype (green circle). **c**, A multiple alignment of orthologous sequences of His3 for segment 2 for which we incorporated almost all extant amino acid states from 21 yeast species (blue bars) and 10-100% extant states from a set of 396 orthologues (grey bars). **d**, The predicted structure of His3p with amino acid residues that were substituted in our library.

## Estimating fitness of evolutionary-relevant genotypes

We studied His3, a gene coding for imidazoleglycerol-phosphate dehydratase (IGPD, His3p), an enzyme essential for histidine synthesis. In a multiple alignment of His3 orthologues from 21 yeast species we identified 686 extant amino acid states (**Supplementary Information 1**), which were evenly distributed across the His3p structure (**Fig. 1d**). These 686 substitutions, which occurred over the course of ~400 million years of evolution^30^ (**Fig. 1b,c**) correspond to ~10^83^ sequences, even a tiny fraction of which would be too many to survey. Thus, we sectioned His3 into 12 independent segments such that the full combinatorial set of substitutions that have occurred in His3 during yeast evolution comprised 10,000-100,000 genotypes per segment (see Methods and **Supplementary Fig. 1a**). The 12 segments were of similar length, constrained by the molecular methods employed for library construction (see Methods), and covered a diverse range of secondary structures and functional elements (**Supplementary Fig. 1c**). For each of the 12 segments of His3 we performed an independent experiment surveying its fitness landscape. For each segment we used degenerate oligonucleotides to construct genotypes consisting of combinations of amino acids present in extant His3 sequences, and determined the fitness conferred by these genotypes by expressing them in a *Δhis3* strain of *S. cerevisiae* and measuring the rate of growth **(Supplementary Fig. 1b)**. This way we assayed the fitness landscape of the genotype space that was traversed over the course of the last ~400 million years of evolution^30^.

Across 11 experiments we measured fitness for a total of 4,018,105 genotypes (875,151 unique amino acid sequences) with high accuracy. Of these, 422,717 consist solely of combinations of extant amino acid states from His3 orthologues, while the remaining genotypes incorporate other amino acid substitutions **(Supplementary Table 1** and Methods). For one segment, 9, the accuracy of our experiment was low, and it was not used in cumulative analyses. For each segment we measured fitness for 60% - 99.8% of all possible genotypes from the combinatorial set of selected extant amino acid states found in 21 yeast species and a smaller fraction of combinations found across all domains of life (**Supplementary Table 1**), characterizing the evolutionary relevant fitness landscape **(Fig. 1b)**. For segment 3 for instance, 11 out of 17 amino acid sites had more than one extant amino acid state: L145=2, L147=2, Q148=3, K151=2, V152=2, D154=3, L164=3, E165=4, A168=2, E169=4, A170=4, with the full yeast combinatorial set consisting of 2*2*3*2*2*3*3*4*2*4*4 = 55,296 genotypes out of which we determined the fitness for 48,198, or 87% of the possible yeast extant states combinations in our library.

A substantial proportion of combinations of extant amino acid states led to genotypes with low fitness (**Fig. 2, Fig. 3a,b, Supplementary Fig. 2**), an observation that takes into account the false discovery rate in our data (**Supplementary Table 1**). This observation could be explained by i) some extant amino acids having a universally deleterious effect, ii) some amino acid states exerting a negative effect on fitness because of intergenic interactions with other *S. cerevisiae* genes, or iii) by epistatic interactions between the extant amino acid states within His3^31^. We exclude the possibility that some extant amino acid states had a universally deleterious effect because no extant amino acid states were present only in unfit genetic backgrounds, genotypes conferring a fitness of zero (**Fig. 3c**). We exclude the possibility that some extant amino acid states disrupt intergenic interactions because the complete His3 coding sequences from extant species fully complemented a His3 deletion in *S. cerevisiae* (**Supplementary Fig. 3c**). Thus, the observed genotypes with low fitness can only be explained by epistatic interactions among extant amino acid states within the His3 gene in the same or different segments. Remarkably, 85% (330/389) of substitutions between extant amino acid states had substantially different effects on fitness in different backgrounds (**Fig. 3d**). By contrast, only 15% of amino acid substitutions that occurred in His3 evolution are truly neutral, in the sense that they do not exert strong influence on fitness in any genetic background. Three quarters of the universally neutral substitutions were observed in the disordered region of the protein (44/59). Thus, the His3 fitness landscape across the 11 segments with high accuracy was strongly influenced by epistasis on a macroevolutionary scale, i.e. the impact of an extant amino acid state on fitness often depends on the background in which it occurs^31–34^. An epistatic fitness landscape is rugged in the sense that evolving genotypes must avoid fitness valleys that emerge through deleterious combinations of amino acid states that may also be found in fit genotypes^18,19,34–36^. Characterizing the ruggedness and the mechanisms that determine the underlying epistasis becomes the primary challenge in understanding the fitness landscape of His3.

**Figure 2.**
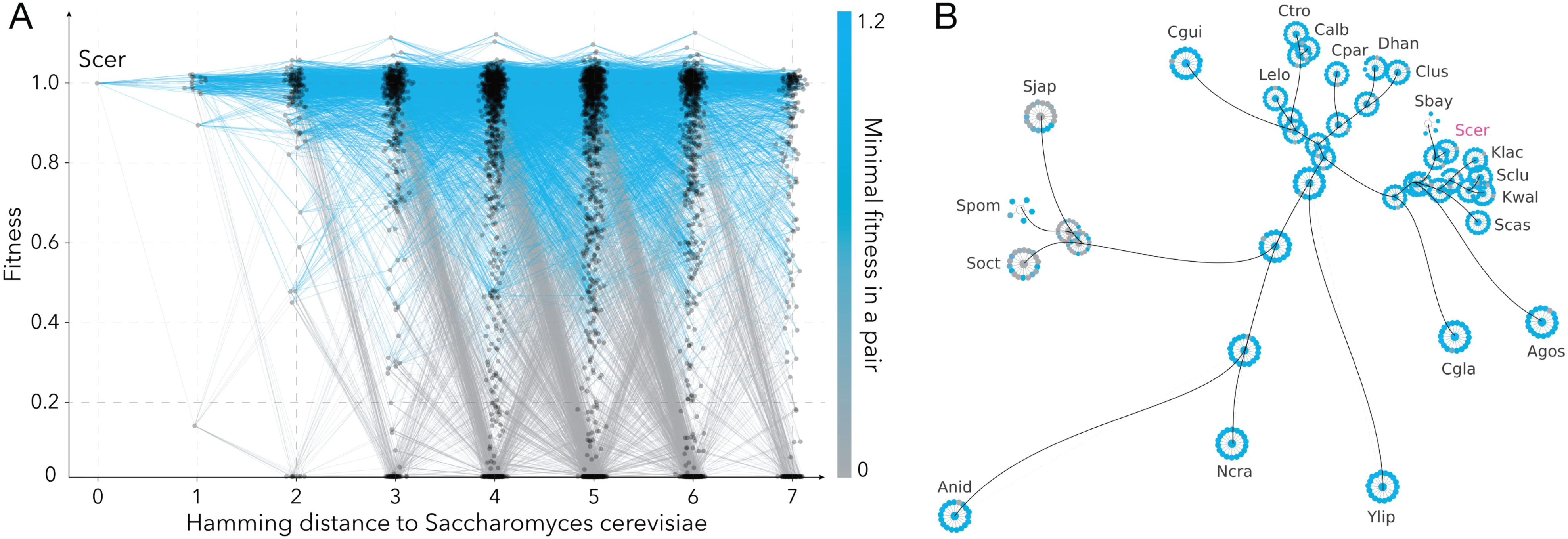
Visual representations of the fitness landscape. **a,** The fitness landscape for all assayed genotypes in segment 7. Nodes represent unique amino acid sequences with edges connecting those separated by a single amino acid substitution. Colour saturation represents the minimum fitness of the two connected nodes. **b,** For segment 7, fitness of ancestral and extant nodes and genotypes one substitution away from the nodes in the background of *S. cerevisiae* gene on the yeast phylogeny (black lines), are shown in colour ranging from grey (lowest fitness) to blue (highest fitness).

**Figure 3.**
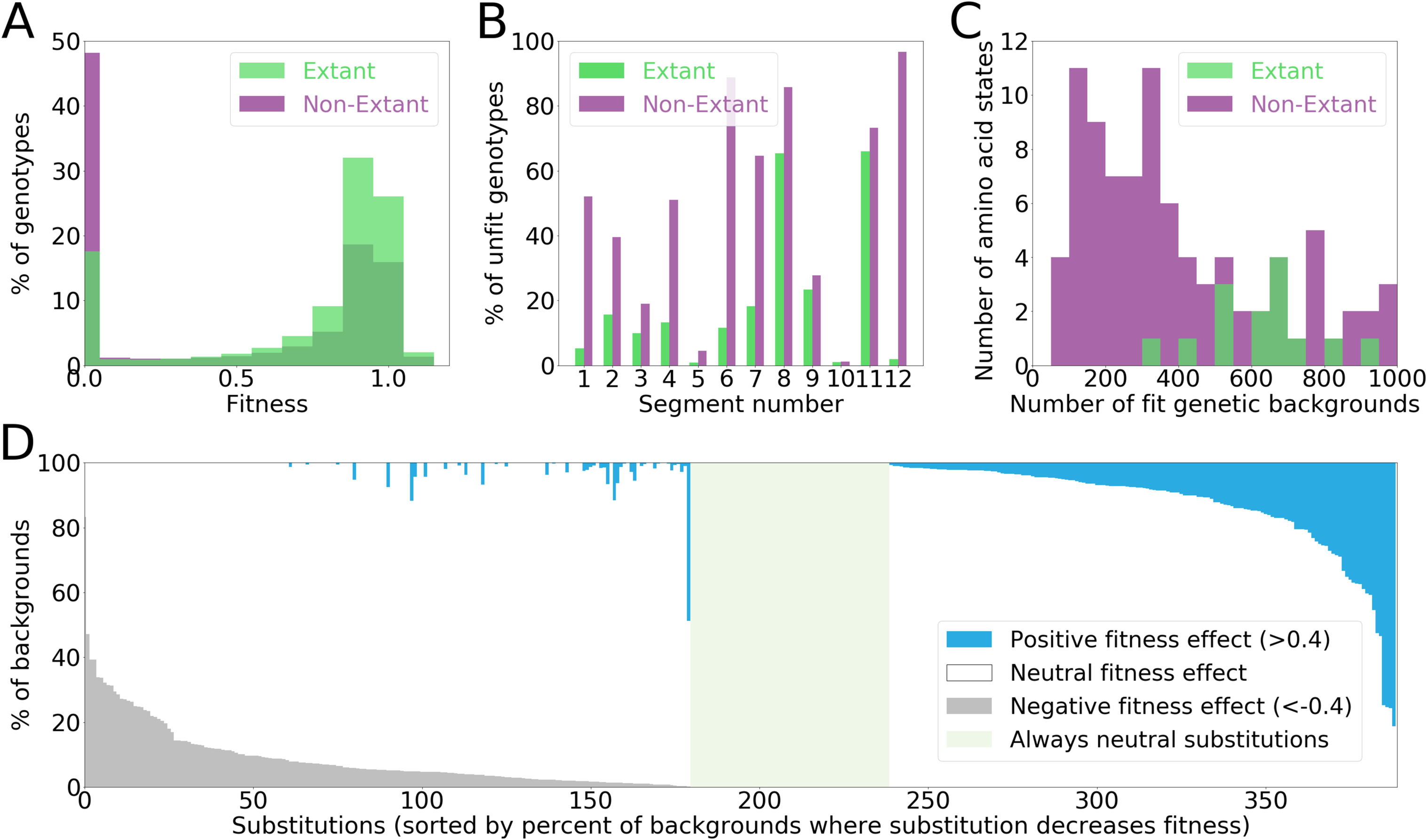
Fitness distributions. **a,** The distribution of fitness for genotypes composed of combination of extant amino acid states (green) and non-extant amino acid states (purple) at the same positions. **b**, The fraction of unfit genotypes per segment among genotypes consisting entirely from extant amino acid states (green) and those incorporating non-extant amino acid states (purple). **c**, The number of genotypes with high fitness that incorporates specific amino acid states. For each amino acid state, the number of genetic backgrounds that contain that amino acid state and are fit (fitness > 0) are shown. **d**, The percent of backgrounds in which a specific substitution is neutral (white), beneficial (blue) or deleterious (grey). The region marked in green shows substitutions that never have large effects (> 0.4) on fitness. Beneficial and deleterious effects are shown only if the frequency for a given substitution was higher than the false discovery rate (**Supplementary Table 1**). Data from segment 9 were excluded for this figure.

## Unidimensional epistasis of the His3 fitness landscape

The ruggedness of the fitness landscape can be characterized by different measures of complexity of the underlying epistatic interactions. In the simplest case, epistasis may be unidimensional, in the sense that the fitness landscape can be described as a function of an intermediate variable, the fitness potential^37–39^. The fitness landscape is a function from the space of genotypes to fitness. In analogy with a scalar field, we can characterize the ruggedness of this function with standard measures of complexity if genotypes are arranged in a linear space. The simplest case is that of a linear predictor called the fitness potential: *p* = c_1_*x*_*1*_ + *c*_*1*_*x*_*2*_ + … + c_n_*x*_*n*_, where c_i_ is a coefficient and *x*_*i*_ is a binary variable that signifies the presence (1) or absence (0) of a given amino acid at a given position. By definition, *e*^*p*^ describes a non-epistatic fitness landscape because the effect of every substitution is multiplicative and it depends only on the associated c. Any other function of *p* leads to epistasis. If the *f(p)* function is “simple”, meaning that is has a small number of local extremas, such as a bimodal function, the epistasis is called unidimensional^39^. The limitation of simplicity of *f(p)* is necessary because any function f_0_(x_1_, …, x_n_) can be represented by a function *f’(p)* and choosing appropriate coefficients c_1_, …, c_n_ in *p*. Thus, a simple *f(p)* leads to unidimensional epistasis because the entire genotype space can be reduced to a single dimension^39^.

To quantitatively determine how well fitness differences between genotypes can be explained by unidimensional epistasis we used a deep learning approach to estimate the coefficients c for each allele x in the fitness potential and determine the best unidimensional function of *p* that best approximated the fitness landscape. We used a dense neural network architecture composed of three layers. Each neuron in the architecture performed a linear transformation of its input and then applied a nonlinear (sigmoid) function. Hence, by using one neuron in the first layer we obtained a linear combination of the contributions of each amino acid state to fitness potential, which was then non-linearly mapped to fitness by the three layers of the neural network architecture (see Methods; **Supplementary Fig.4**). Ten segments were described by a threshold function in which organismal fitness remains constant with decreasing fitness potential and then is rapidly reduced to lethal after a certain threshold (**Fig. 4a**). The ability of the cliff-like threshold fitness function^40^ to predict fitness from genotype varied between the His3 segments from near perfect (r^2^=0.97) in segment 7, to relatively poor (r^2^=0.44) in segment 5 (**Supplementary Fig. 5**). Thus, while the fitness landscape of His3 is approximately unidimensional for some segments, it has a higher degree of complexity for others.

**Figure 4.**
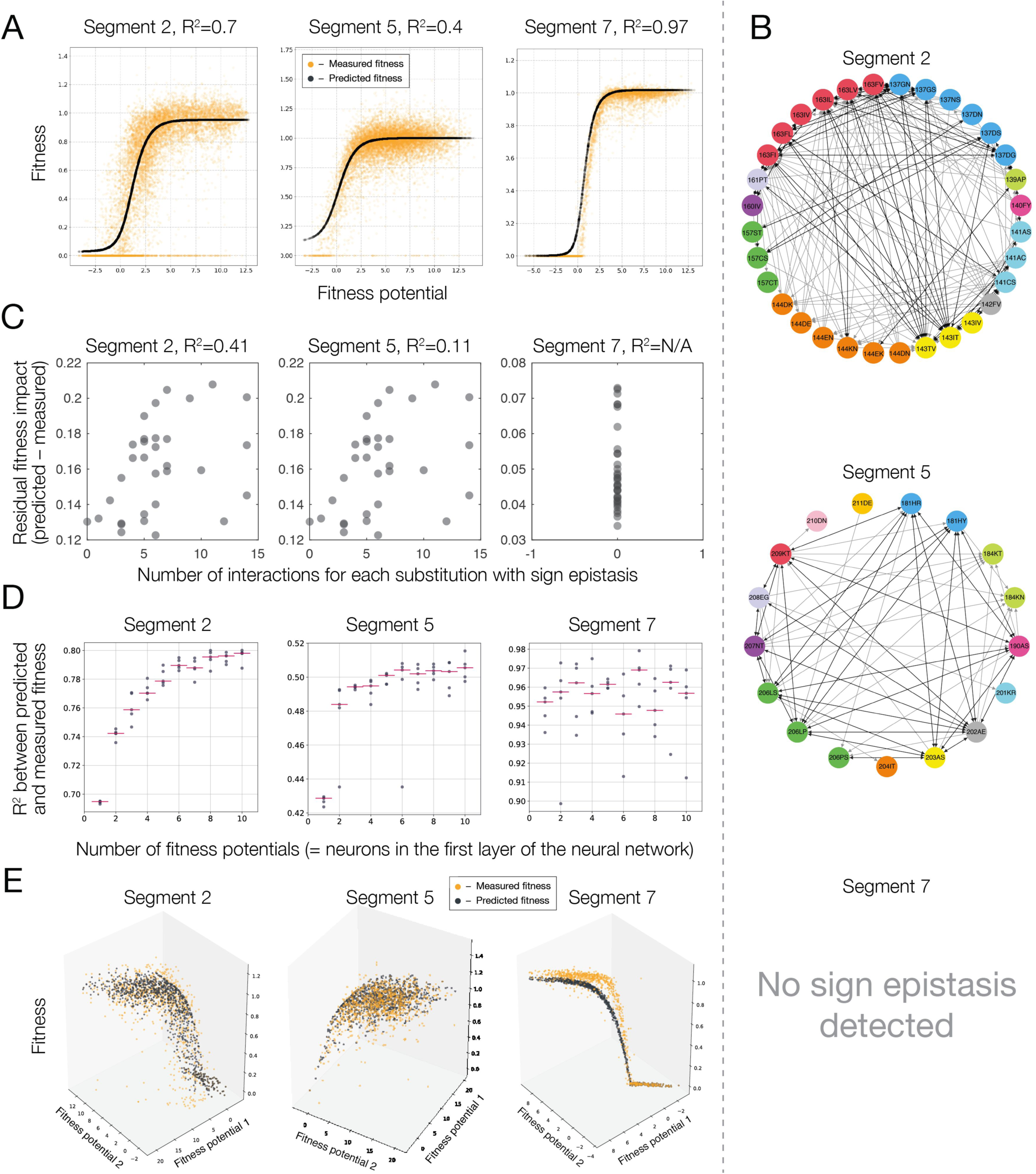
Epistasis and the His3 fitness landscape for segments 2, 5 and 7. **a**, Fitness as a function of a single fitness potential (black curve, the fitness of individual genotypes is orange). **b**, A network depiction of sign epistasis between amino acid substitutions. Colour coded sites with reciprocal sign epistasis (black lines) and unidirectional interactions (grey arrows) are shown. **c**, Genotypes containing substitutions with a higher number of sign epistatic interactions are less likely to be fit by the threshold function of the fitness potential. **d**, Increasing the number of neurons in the first layers of the neural network, which is equivalent to increasing the number of underlying fitness potentials, leads to more accurate models for segments with detected sign epistasis. Each dot corresponds to an independent optimization of model parameters. **e**, Fitness as a function of two fitness potentials (black dots, measured fitness is depicted in orange).

## Ruggedness and multidimensional epistasis of the His3 fitness landscape

Ruggedness is a general property of fitness landscapes that quantifies the accessible paths of high fitness that connect fit genotypes^41–43^. A path between highly fit genotypes is inaccessible when one of the intermediate genotypes has low fitness^6,20–22,41,44^ *(e.g.* for genotypes AB and ab, the intermediate are aB and Ab). Such instances also manifest in sign epistasis on the fitness landscape, that the same substitution may be beneficial or deleterious when occurring in a different genetic background^44,45^. To quantify the ruggedness of the His3 fitness landscape we identified instances of sign epistasis: substitutions between extant amino acid states that were strongly beneficial or strongly deleterious (change in fitness of > 0.4 in absolute value) depending on the background in which they occurred^44^. Some of these instances may be due to miscalled fitness of very few genotypes. Therefore, we considered a pair of extant amino acid states to be under sign epistasis only when sign epistasis was observed in a statistically significant number of different genetic backgrounds (see Methods).

An example of sign epistasis is the substitution C141S in the second segment that had an opposite effect on fitness depending on amino acid at site 143 (I, V or T). The substitution I143T in turn exhibits sign epistasis depending on the amino acid at site 163 (F, I, V or L) **(Fig. 5a)**. These epistatic interactions can be represented by a graph in which nodes represent a pair of extant amino acid states at a specific site and nodes are connected by edges if strong sign epistasis has been detected between them (C141S - I143T - I163F) **(Fig. 4b)**. We found that 86 out of 128 (67%) sites in our library exhibit sign epistasis and 46% (59/128) exhibit reciprocal sign epistasis. Most sites showed a sign epistatic interaction with multiple other sites (**Fig. 5c,d**) demonstrating that, although sign epistasis affects few genotypes, it leads to a fitness landscape that requires the interaction of multiple sites for proper characterization.

**Figure 5.**
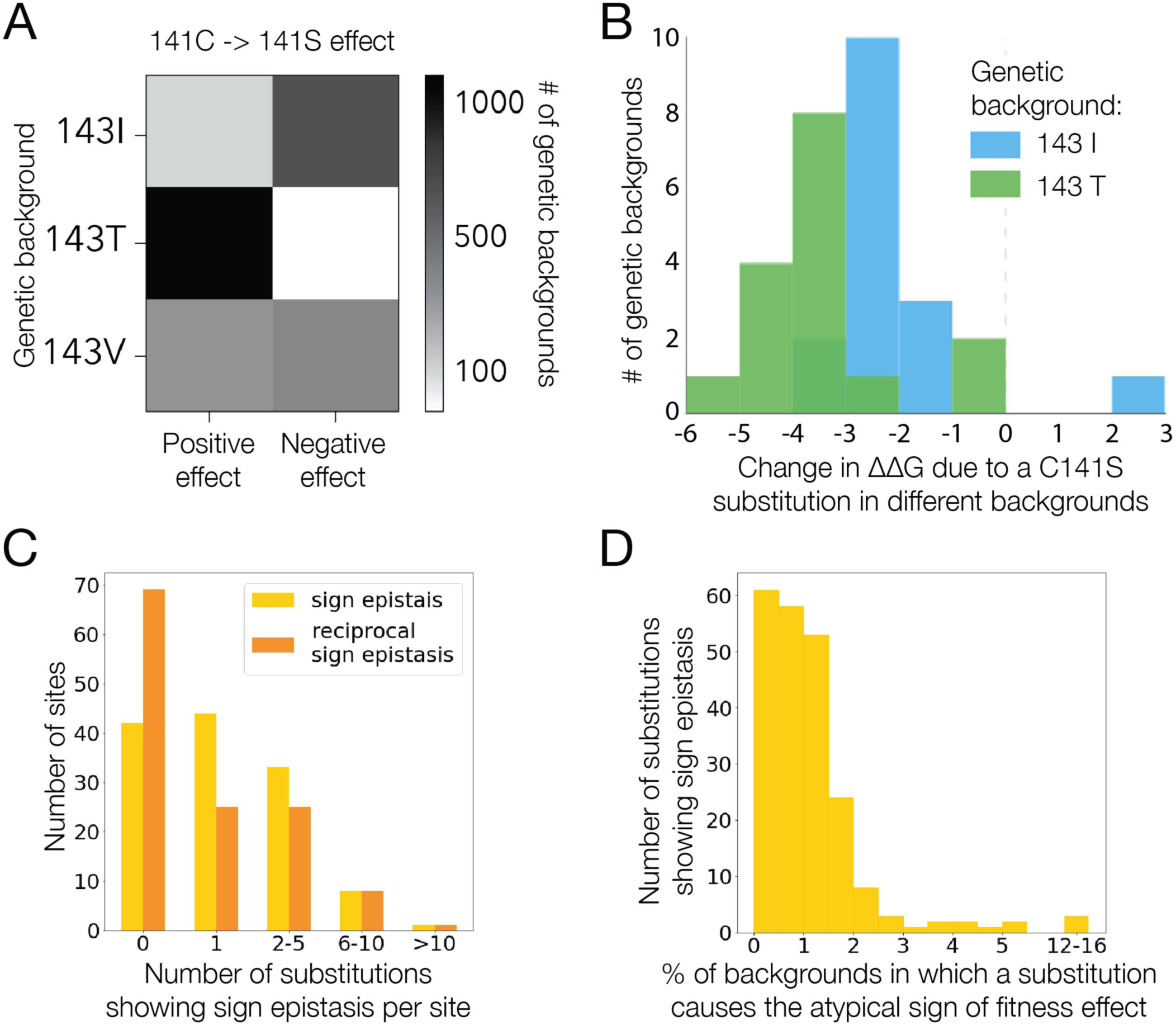
Sign epistasis. **a**, Substitution C->S at site 141 in segment 2 more frequently has a positive effect on fitness in the background of T at site 143, a negative effect in the background of 143I and is equally likely to be strongly deleterious or strongly beneficial in the background of 143V. **b**, Predicted change in free energy following a C141S substitution in all genetic backgrounds with an I or T at 143 and that are closer than four mutations away from *S. cerevisiae.* **c**, Distribution of the number of substitutions at each site under sign (yellow) and reciprocal sign (orange) epistasis. Sites with 0 interactions do not exhibit sign epistasis. **d**, The fraction of genotypes in which the substitution under sign epistasis has the less frequent effect on fitness.

The complexity of interaction of sites can be estimated by using the graph of sign epistasis where vertices represent a substitution and edges connect vertices with sign epistasis between them. If only few substitutions display sign epistasis then such a graph would signify that the fitness landscape is relatively smooth, alternatively, a highly-interconnected graph of such interactions signifies a more rugged landscape^4,5,41–43^. To measure the relative fitness complexity, we used the maximum clique size of a graph, which approximates the maximum number of simultaneously interacting substitutions. In our data, this measure ranged from two to seven depending on the segment (**Supplementary Fig. 6**). The ruggedness of the fitness landscape of His3 is high for most segments, such as segment 5, where it is necessary to consider the simultaneous interaction of at least seven sites to accurately predict the fitness of genotypes consisting of extant amino acid states at these sites^38,40^.

Sign epistasis can appear when fitness is described by a unidimensional function of the fitness potential, for example, when the fitness landscape is a unimodal function with an optimum in an intermediate range of the fitness potential^41,45^. However, sign epistasis may also be a sign of multidimensional epistasis, when a unidimensional function of the fitness potential cannot fully describe genotype fitness^39^. Many genotypes were predicted poorly by a unidimensional function of the fitness potential (**Supplementary Fig. 5a**). Two lines of evidence suggest that such genotypes reveal the presence of multidimensional epistasis. First, genotypes with a higher number of substitutions influenced by sign epistasis were less well-predicted by a unidimensional fitness function (**Fig. 4c and Supplementary Fig. 7b**). Second, we explain a larger fraction of genotypes by using a more complex neural network architecture accommodating multiple fitness potentials instead of one. We found that increasing the amount of neurons in the first layer of the neural network architecture, which is equivalent to increasing the number of independent fitness potentials, gradually improves the prediction power of the obtained models for most of the segments (**Fig. 4d**). Thus, adding dimensions to the function of fitness potential increases the prediction power of the model. For example, for a two-dimensional case fitness was described by *f_1_(p_1_,p_2_)* with *p*_*1*_ = *a*_*1*_*x*_*1*_ + a_1_*x*_*2*_ … a_n_*x*_*n*_ and *p*_*2*_ = *b*_*1*_*x*_*1*_ + b_1_x_2_ … b_n_*x*_*n*_. For several His3 segments, a fitness function with multiple underlying fitness potentials described the fitness landscape more accurately than a simple unidimensional function of a single fitness potential (**Supplementary Fig. 7a**). For instance, for these segments, fitness function of two fitness potentials described the shape with a higher degree of accuracy than a function of a single fitness potential (**Fig. 4d,e**). By contrast, epistasis in segment 7 is entirely unidimensional (**Fig. 4d,e and Supplementary Information 2**); we do not see any improvement in the model’s predictive power when adding extra dimensions.

## Evolutionary trajectories on the His3 fitness landscape

On a smooth fitness landscape, evolution can proceed along any of the evolutionary paths connecting two fit genotypes, as none of the intermediate genotypes confer low fitness (see Box 2 in [46]). Alternatively, the fitness landscape is rugged when it contains non-connected fitness peaks, such that there are no viable paths between some pairs of genotypes that confer high fitness^4,5^. In other words, the presence of deleterious intermediate genotypes between highly fit ones leads to inaccessibility of some evolutionary trajectories between extant or ancestral sequences^6,20–22,44^. The simplest explanation for the substantial ruggedness of the landscape observed in many of the His3 segments lies in the unidimensional threshold fitness function (**Fig. 6a**). On a threshold function a combinations of substitutions, all of which are neutral in some genetic backgrounds, can take a genotype beyond the fitness threshold through their additive effect on fitness potential, making some genotypes inaccessible for evolution (**Fig. 6a**). Between any two fit genotypes, the fraction of intermediate genotypes that are unfit depends on the fitness potential of the two parental genotypes (**Fig. 6b**). Evolution between two fit genotypes with high fitness potential can proceed unhindered because all intermediate genotypes also have high fitness potential and, consequently, high fitness. Conversely, when both fit genotypes are located close to the threshold, many of the intermediate genotypes between them have low fitness and many evolutionary paths between them are inaccessible (**Fig. 6c**). Thus, the cliff-like threshold fitness function is the major determinant of the observation that not all paths between two fit genotypes are accessible to evolution (**Fig. 6b**). We find that unfit intermediate genotypes are in genetic proximity with each other and are on a limited number of paths; the fraction of inaccessible paths is smaller than if the same number of unfit genotypes were distributed randomly in genotype space (**Fig. 6d,e**).

**Figure 6.**
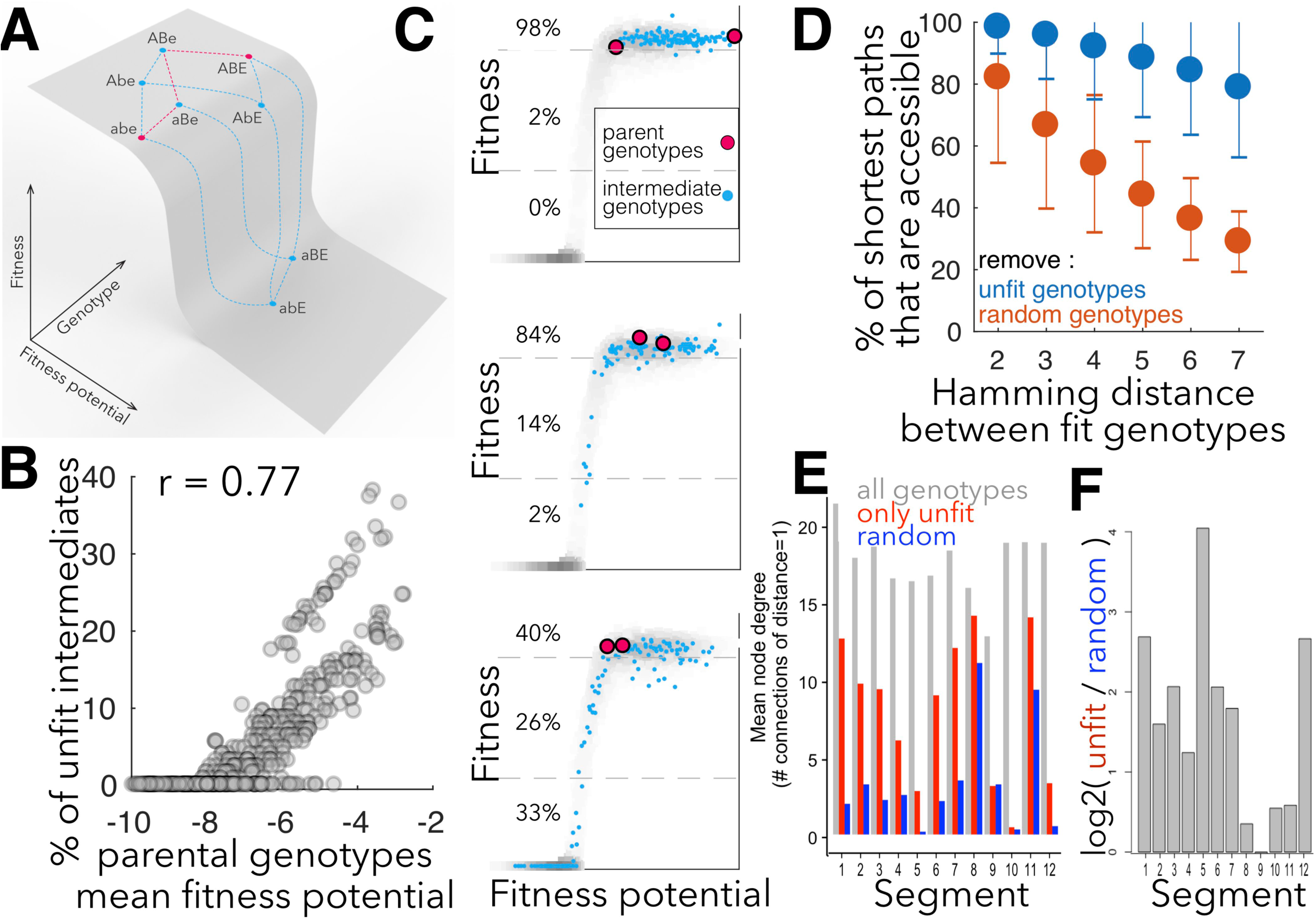
Analysis of evolutionary pathway accessibility. **a,** A threshold fitness potential function can lead to some paths being inaccessible between two genotypes of high fitness (abe, ABE) if the joint contribution of several alleles to the fitness potential (abE, aBE) leads to the fitness potential below the threshold. **b,** The fraction of unfit intermediate genotypes between two fit genotypes as a function of their average fitness potential. **c**, The grey area represents all genotypes in segment 7. When two fit genotypes (red dots) have high fitness potential, many paths between them will be accessible because many intermediate genotypes will also have high fitness potential and fitness (blue dots). **d**, The fraction of accessible shortest paths between two fit genotypes with unfit genotypes from data (orange) or the same number of randomly selected genotypes (grey), shown as a function of Hamming distance between two fit genotypes. Error bars are standard deviation. **e**, On a graph with genotypes connected by edges if they are one substitution apart we calculate the degree of connectivity (number of edges for each node) for all genotypes (blue), only unfit (fitness = 0) genotypes (orange) and a graph with the same number of nodes as the graph with unfit genotypes but with nodes chosen at random (grey).

The effect of synergistic epistasis dominates the His3 fitness landscape, affecting over 85% of amino acid substitutions from our library that occurred in His3 evolution. This synergistic epistasis may reflect the free energy of the protein^10,47,48^, as evidenced by a correlation between the fitness potential and the impact of substitutions on the free energy of His3p (**Supplementary Fig. 8**). Similarly, instances of sign epistasis may also be explained by changes in protein stability; for example, in the 143T background C141S increased fitness and also had a positive effect on stability (**Fig. 5b)**. Consistent with protein stability contributing to the observed sign epistasis we find that sites that exhibited reciprocal sign epistasis are close together in the His3p structure **(Supplementary Fig. 8)**. However, an additive contribution to free energy can lead only to a unidimensional fitness function^47^, indicating that other non-additive mechanisms, such as catalytic activity or inter-subunit interactions, or a non-additive model of free energy, must be responsible for the multidimensionality of the His3 fitness landscape.

## Inference of inter-segmental epistatic interactions in the His3 gene sequence

Epistasis may be caused by interaction among positions within a segment (intra-segmental epistasis) or by interaction of the segment with the rest of the *S. cerevisiae* His3 sequence (inter-segmental epistasis). The contribution of inter- versus intra-segmental interactions can be decoupled. Given two fit genotypes (e.g. ABC & abc in one His3 segment), any unfit intermediate states (e.g. aBc in the same His3 segment) must be due to intra-segmental epistasis because the rest of the protein remains constant. For each segment we took as a measurement of intra-segmental epistasis all pairs of fit genotypes and calculated the proportion of unfit intermediate genotypes as a function of the Hamming distance between the two fit genotypes. We then compared this proportion with the total proportion of all unfit genotypes as a function of Hamming distance from *S. cerevisiae*, a measurement that includes both inter- and intra-segmental epistasis. We found three times more inter-segmental than intra-segmental epistasis **(Supplementary Fig. 9)**, likely because a single segment provides a much smaller target space for interactions than the entire His3 protein. The proportion of sites under epistatic interactions increased exponentially with Hamming distance (**Supplementary Fig. 9**), analogous to Orr’s snowball, the accumulation of genetic incompatibilities in the course of speciation^31,49^.

## Conclusions

The concept of the fitness landscape introduced by Sewall Wright (Figures 1 and 2 in [6]) is an indispensable tool for understanding multiple biological phenomena^1–5^. Experimental high-throughput assays of random mutations have begun to unravel some local properties of fitness landscapes^4^. Here, we described a fitness landscape on a macroevolutionary scale by focusing on amino acid states that have been put through the sieve of natural selection. We found that only 15% of substitutions that fixed in the evolution of His3 are universally neutral. For the remaining 85%, substitutions from His3 evolution had a profound influence on each other’s effect on fitness, providing an experimental confirmation that epistasis is one of the defining features of molecular evolution^33^. Substitutions that occur in evolution have properties vastly different from those of random mutations, which are mostly deleterious^23^. Therefore, the way in which combinations of extant amino acid states affect fitness may also be different from that of combinations of random mutations. Unexpectedly, we found that the interaction of extant amino acid states was dominated by synergistic epistasis in a manner similar to that previously found for random mutations^7–10,16^. However, the accumulation of random mutations leads to low fitness much faster than the accumulation of extant amino acid states (compare Figure 2 from [16] and Figure 3b from [10] to **Fig. 2a**).

The experimental data showing that 85% of amino acid states found in extant species confer low fitness in a different genetic background lends strong support to the notion that epistasis is a key factor in protein evolution^31,33^. We showed that the fitness landscape of several segments of the His3 gene cannot be reduced to a single unidimensional forms of epistasis, with a function of multiple fitness potentials providing a more accurate description of the fitness landscape. By contrast, large-scale fitness landscapes incorporating multiple random mutations away from the wildtype sequence in a constant test environment have not displayed evidence of multidimensional epistasis^8-10,16^; however, it appears to be a more prevalent factor among substitutions that have been subject to positive selection ^12–16,18–22,34–36^. We also found that up to 67% of sites with an extant amino acid state were influenced by sign epistasis, resulting in a rugged fitness landscape and a limited number of fitness ridges connecting extant sequences for most His3 segments. Overall, the evolutionary-relevant section of the His3 fitness landscape is best described as a fitness ridge, with the crest of the ridge defined by a fitness potential. In some cases, the crest is multidimensional requiring several independent underlying fitness potentials. Evolution can proceed unhindered along the crest **(Fig. 6a,c)**, however, pathway availability declines rapidly when evolution proceeds close to the edge of the fitness ridge.

## Acknowledgements

We thank Ben Lehner for a thorough study of the concept behind our experiments and Julia Domingo for technical assistance with preliminary analysis of our data. Jochen Hecht of the CRG Genomics Unit and Svetlana Belogorodtdeva and Roman Belogorodtsev provided technical assistance. The work was supported by HHMI International Early Career Scientist Program (55007424), the MINECO (BFU2012-31329, BFU2012-37168, BFU2015-68351-P and BFU2015-68723-P), Spanish Ministry of Economy and Competitiveness Centro de Excelencia Severo Ochoa 2013-2017 grant (SEV-2012-0208), the Unidad de Excelencia María de Maeztu funded by the MINECO (MDM-2014-0370), Secretaria d’Universitats i Recerca del Departament d’Economia i Coneixement de la Generalitat AGAUR program (2014 SGR 0974), the CERCA Programme of the Generalitat de Catalunya, Russian Foundation for Basic Research grant (18-04-01173), the European Union’s Horizon 2020 research and innovation programme under the Marie Skłodowska-Curie programme (665385) and the European Research Council under the European Union’s Seventh Framework Programme (FP7/2007-2013, ERC grant agreement 335980_EinME and Synergy Grant 609989).

## Author contribution

ISP, LBC and FAK conceived the general approach of the study. FAK, LBC, VOP, LE and GJF participated in detailed experimental design. VOP, DRU and EVP spearheaded the experiment, data analysis and main data interpretation, respectively. LE performed a large fraction of the experimental work. AVA and SYaA participated in working on dimensionality. KSS, ASM, NSB, DNI and GJF participated in data analysis. LBC and FAK wrote the draft.

## Methods

### Data access

The raw and processed data have been submitted to the NCBI Gene Expression Omnibus (GEO;http://www.ncbi.nlm.nih.gov/geo/) under accession number GSE99990. A virtual machine containing a running version of the data processing pipeline is available as a Docker image https://hub.docker.eom/r/gui11aume/epi/. The scripts to reproduce the figures are on Github at https://github.com/Lcarey/HIS3InterspeciesEpistasis.

### Study design

The His3 gene was selected for three principal reasons, it is short, conditionally essential and was not known to be involved in protein-protein interactions. Studying 20^220^ variants of His3 is impossible, thus, we have chosen an approach to survey the fitness landscape in a manner that would elucidate the area most relevant to His3 evolution while managing the technical limitations of our experimental design. We considered amino acid states found in extant species, focusing on yeast species, which translated into a full combinatorial set of ~10^83^ unique genotypes. Technically it is feasible to measure fitness of up to 100,000 unique genotypes in a single growth experiment. Therefore, we split the His3 gene into 12 independent segments such that the full combinatorial set of extant amino acid states from 21 yeast species in each segment was 10,000 - 100,000 genotypes. We then considered the combinatorial library for each segment in an independent growth experiment, which allowed us to study a tractable section of the sequence space while considering trajectories across a vast part of the space connecting extant species (**Fig. 1a**). We constructed these combinations in 12 plasmid libraries and transformed them into a haploid His3 knockout *S. cerevisiae* strain. Growth rate (fitness) of yeast carrying different mutations in His3 was measured using serial batch culture in the absence of histidine.

We split the His3 gene sequence into segments in a manner agnostic to the structure of the His3 protein (**Supplementary Fig. 1a**). For technical reasons, a segment consisted of two variable regions with a constant region between them (**Supplementary Fig. 1b**). All growth experiments were performed independently for each segment, with the exception of one experiment on a limited group of genotypes from each segment which was done for the normalization of fitness values across different segments (**Supplementary Fig. 3**).

As a control, we measured the rate of growth of *S. cerevisiae* whose entire His3 gene sequence came from another distant species. We found that the replacement of an entire gene sequence of His3 leads to wild-type rates of growth of *S. cerevisiae* even when the His3 sequence comes from very distant yeasts, as far as *S. pombe* (**Supplementary Fig. 3**). Therefore, His3 appears to be an independent unit of the fitness landscape and is a good model for the study of fitness landscapes of an isolated gene.

### Measuring fitness

#### Plasmid construction

The His3 open reading frame of *S. cerevisiae* was PCR amplified with its regulatory region from 622 base pairs (bp) upstream of the open reading frame (ORF) to 237 bp downstream of the ORF, using primers 126 and 127 (see **Supplementary Table 1**) from the wild-type prototroph strain FY4. The PCR product was cloned into vector pRS416 using Gibson assembly (NEB, E2611S). The His3 orthologues from other species were amplified from genomic DNA using designed primers (**Supplementary Table 1**) and were cloned into the vector pRS416_his3, replacing the ORF of *S. cerevisiae* by Gibson assembly (NEB, E2611S). Since the His3 orthologue from *A. nidulans* contains an intron, the whole open reading frame was initially cloned into the vector, and the intron was later removed by PCR-amplifying the whole plasmid without this sequence, followed by recircularization.

#### Genomic DNA extraction

Genomic DNA from fungi *(Saccharomyces cerevisiae, Saccharomyces bayanus, Candida glabrata, Saccharomyces castellii, Kluyveromyces lactis, Eremotheciumgossypii, Debaryomyces hansenii, Lodderomycese longosporus, Aspergillus nidulans, Schizosaccharomyces pombe, Candida guilliermondii, Saccharomyces kluyveri, Kluyveromyces waltii)* was extracted using MasterPure™ Yeast DNA Purification Kit according to the manufacturer’s instructions (Epicentre, MPY80200).

#### Mutant library construction

Twelve independent mutant libraries, each for different regions of His3 **(Supplementary Table 1)**, were generated based on the results of multiple alignment of His3 orthologues. The alignment was built using the ClustalW alignment feature of the MEGA 6.0 software package^50^ and user-corrected.

Mutant libraries were constructed by fusion-PCR, leaving two variable regions separated by a constant region. For each library, two contiguous fragments of His3 were amplified independently, using 1 μg of *S. cerevisiae* (strain FY4) genomic DNA in separate Phusion polymerase reaction mixes (Thermo Fisher Scientific, F530S) in GC buffer. For each PCR, one of the primers was a degenerate oligonucleotide with a constant part at the 5’ end required for the fusion-PCR; the other primer was either 126 or 127. The degenerate primer approach led to the integration of non-extant amino acid sequences due to the redundancy of the genetic code. Consider the amino acid Phe in *S. cerevisiae* coded by the codon TTT. When incorporating an extant orthologous state Trp (TGG) two independent T -> G nucleotide mutations will be incorporated creating the codons TTG (Leu) and TGT (Cys). If these two amino acids were not found in other species then they would be non-extant. The cycling conditions for the PCR were 98°C for 30 s; 98°C for 20 s, 60 °C for 30s and 72°C for 1 min (25 cycles); and 72°C for 5 min. The products were column-purified (QIAGEN, QIAquick PCR purification kit, 28104), eluted in 50 μl and mixed in equimolar proportion. The fusion-PCR was carried out by diluting 10 μl of the mix to 25 μL of standard Phusion polymerase reaction mix in GC buffer. The cycling conditions of the fusion-PCR were 98°C for 30 s; 98°C for 30 s, 60°C for 2 min and 72°C for 1 min (25 cycles); and 72°C for 5 min. The product of fusion was purified from agarose gel (Qiagen, MinElute Gel Extraction Kit, 28604) and eluted in 10 μl of water. 10 μl of the product was used as a template for additional 5 cycles of PCR reaction in Phusion polymerase reaction mix (Thermo Fisher Scientific, F530S) in GC buffer, using primers 126 and 127. The cycling conditions were as follows: 98°C for 30 s; 98°C for 20 s, 60°C for 30 s and 72°C for 1 min (5 cycles); and 72°C for 5 min. The product was column-purified (QIAGEN, QIAquick PCR purification kit, 28104), and used as an insert for Gibson assembly.

To create a library of His3 mutants, pRS416 plasmid was amplified using primers 128 and 129. The insert was cloned into the vector using Gibson assembly (NEB, E2611S). Ligated products (200 - 300 ng/μL) were desalted by drop dialysis using 13 mm diameter, Type-VS Millipore membrane (Merck Millipore, VSWP01300). 20 μL ElectroMAX DH10B competent cells (Invitrogen, 18290015) were electroporated with 3 μL ligated products. 0.01% of the electroporated bacteria were plated on ampicillin-containing medium in order to estimate the complexity of the library; the remaining culture was grown overnight in 100 ml of liquid medium, and the plasmid was extracted the next day. For each library, the maximum number of protein sequences that can be generated was computed. Libraries were generated until to total complexity reached at least 3 times this value.

#### Yeast transformation and yeast library generation

For each segment, yeast strain LBCY47 (his3:KanMXleu2Δ0 *met15Δ0 ura3Δ0,* derived from BY4741) was transformed with 50 μg of pRS416_His3 mutant library using lithium acetate transformation and plated onto glucose synthetic complete dropout plates lacking uracil. After 40 hours’ growth at 30°C, approximately 0.5 million yeast colonies were scraped off the plates, mixed together and washed 2 times with 100 ml of PBS.

#### Bulk competition

4×10^9^cells were inoculated into 500 ml of glucose synthetic complete dropout medium lacking uracil with 200 mg/L of G418, and grown at 30°C at 220 RPM for 6-8 h in order to eliminate clones with low fitness irrespective of histidine biosynthesis. Cells were later pelleted and washed with 50 ml of PBS. Approximately 10^10^ cells were inoculated into 1 L of synthetic complete dropout medium lacking histidine, and grown at 30°C at 220 RPM for 168 h with 12 h between bottlenecks: ~10^10^ cells were transferred into fresh medium ~10^8^ cells from the culture were kept as sample for the given time point. Bulk competition for each library of mutants was done in two replicates to account for biological variability.

#### NGS library preparation

The relative abundance of yeast mutants was measured in 3 samples: 1) the initial population before selection was applied (t0), 2) the population after 12 h of growth in the selective medium (t1), and 3) the final population after 168 h of growth in the selective medium (t14). In order to extract plasmid DNA, 5×10^9^ cells from each sample were incubated in 300 µL of zymolyase buffer (1 M sorbitol, 0.1 M sodium acetate, 60mM EDTA (pH 7.0), 2 mg/ml zymolyase, 1% 2-Mercaptoethanol) at 37°C for 3 h. The plasmid DNA was purified from the obtained spheroplasts using QIAprep Spin Miniprep Kit (QIAGEN, 27104) according to the manufacturer’s protocol. The obtained DNA was used as a template in a 25 µL of Q5 DNA polymerase reaction mix (NEB, M0491S), using staggered primers for demultiplexing in the following cycling conditions: 98°C for 30s; 98°C for 10s, 60°C for 30s and 72°C for 30s (18 cycles); and 72°C for 2 min. PCR products were purified using Agencourt AM Pure XP beads (Beckman Coulter, A63880), and eluted in 40 µL of TE buffer (pH 8.0). DNA extraction and PCR-amplification were repeated twice for every sample to account for the technical variability.

NGS libraries were prepared from 100 ng of the purified DNA amplicons using Ovation Rapid DR System (Nugen, 0319-32) according to manufacturer’s instructions. Each library was visualized on a Bioanalyzer (Agilent Technologies) and quantified by qPCR with a Kapa Library Quantification Kit (Kapa Biosystems, KK4835). Twelve samples were pooled together (accounting for two biological replicates, two technical replicates and three time points) at the final concentration of 4 nM, and sequenced in the same lane. Samples were sequenced as 125-bp paired-end reads on a HiSeq2500 sequencer (Illumina) with v4 sequencing chemistry.

#### Yeast growth assay

Mutant strains were grown overnight in complete dropout medium lacking uracil. The cultures were diluted to 0.05 OD 600 nm, and grown for 5 h in the same medium. 6 µL of each culture were transferred into 96-well plates in 125 µL of complete dropout medium lacking histidine. Growth of the strains was monitored by measuring OD 600 nm every 10 min using Tecan Infinite M1000 PRO microplate reader equipped with an integrated Stacker module.

The growth rate of individual curves was measured as the inverse of the time to grow from OD = 0.135 = exp(−2) to OD = 0.368 = exp(−1). If the curve did not reach 0.368, the growth was set to 0. Curves that crossed 0.135 or 0.368 were excluded. The growth rate of a clone was measured as the median of 6 independent growth experiments. We excluded from the analysis clones with discordance between growth in solid and liquid medium, clones that could not be sequenced or that showed evidence of contamination by sequencing, and clones such that the Kullback-Leibler divergence of their read counts compared to all synonymous clones was greater than 0.22. The later criterion ensured that the selected clones were not outliers compared to other variants encoding the same protein.

### Growth rates of isolated strains

We isolated 197 strains from all segment libraries of extant amino acid combinations (9-26 strain per segment) and used Sanger sequencing to determine the sequence. For each strain we performed 6 repeats of growth assay and calculated the average growth rate. Fitness values from competition and growth rates are highly correlated (r=0.82 p=10^−48^). Correlation was significant and greater than 0.6 for all segments except S9, where all selected genotypes appeared to be neutral (**Supplementary Fig. 3**).

### Initial data filtering

The individual sequences of the variants were recovered from pair-end reads with the following steps: the constant region between the two variable regions was identified by inexact matching allowing up to 20% errors using the Seeq library version 1.1.2 (https://github.com/ezorita/seeq). The reads are not oriented because the Illumina sequencing adapters were added by ligation, so the constant regions were searched on both reads. Forward and reverse reads were swapped when a match was found on the reverse read. This ensured that all of the sequences are in the same orientation. For multiplexing purposes, the sample identity was encoded in the left and right primers used to PCR-amplify the variants. To demultiplex the reads, we used inexact matching with the candidate primers, allowing up to 20% errors. This approach was faster and less error-prone than using FLASH^51^. To merge the reads, the sequence of the reverse reads was reverse complemented and the constant region was searched by inexact matching allowing up to 20% errors. The position of the constant part in each read indicated how they must be stitched together. In the region of overlap, the consensus sequence was determined by picking the nucleotide with highest quality as indicated in the quality line of the fastq files. If ‘N’ persisted in the final sequence, the reads were discarded. The PCR primers were trimmed so that all the sequences of the same competition would start and end at the same location.

Reads that did not have the constant region, that could not be oriented or that could not be demultiplexed were discarded. The remaining errors in the reads were corrected by sequence clustering. We used starcode version 1.0 [ref. 52] with default parameters and allowing up to two errors. The corrected reads were translated using the genetic code. Variants encoding the same proteins were not merged; they were kept separate for downstream analyses. A running Docker virtual machine with commented scripts to replay the whole the process is available for download at https://hub.docker.com/r/gui11aume/epi/.

### DNA sequence variant frequency calculation and data filtering

The total number of reads for 12 segments, 3 time points and 4 replicas are shown in **Supplementary Table 1**. Genotypes frequencies are defined as the number of reads for a given genotype divided by the total number of reads in that replicate. Mean frequency was calculated over 4 replicas to be used in further analysis. However, to eliminate influence of outliers the median was taken instead of mean if absolute difference between mean and median was greater than the median value. Only genotypes present in both technical replicas of both biological replicas with at least ten reads (summed across all time points) in each of them were kept.

### Noise estimation

The major factors causing noise in genotype frequency measurements are sampling errors, PCR amplification errors and genetic drift during the competition. For all of these factors, the amount of error depends on the genotype frequency. Therefore, we estimated measurement errors as the function of genotype frequency.

For a given segment, time point and a pair of biological or technical replicas for each genotype we calculated the mean frequency and the squared difference of frequencies from these two replicas. We sorted genotypes by mean frequency and grouped them such that each bin contains 5000 genotypes. We calculated the average frequency and the average squared difference in each bin. Additionally, squared error for frequency 0 was set equal to 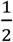. 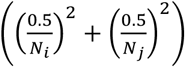, where *N*_*i*_ and *N*_*j*_ are total read numbers in replicas *i* and *j*. Finally, by linear interpolation we obtained dependencies of squared differences as a function of frequency, 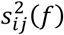,where *i* and *j* are different replicas.

Using squared differences from pairwise comparison of replicas we can estimate variance of mean frequency over four replicas. Let numerate replicas 1, 2, 3, 4 where 1, 2 are technical replicas of the first biological repeat and 3, 4 are the technical replicas of the second biological repeat. Errors coming from the competition (e.g.: genetic drift) are shared for replicas 1, 2 and for replicas 3, 4. Let’s call them Δ*f*_*b*1_ and Δ*f*_*b*2_ and their variances 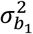 and 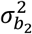 respectively. Technical errors of sampling from the population and from PCR are unique for each replica. Let’s call them Δ*f*_*ti*_,*i* = 1..4 and their variances 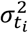, *i* = 1..4 respectively. All variances are function of frequency and writing 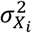 we assume 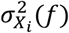.

In the introduced notations the mean frequency over 4 replicas is:

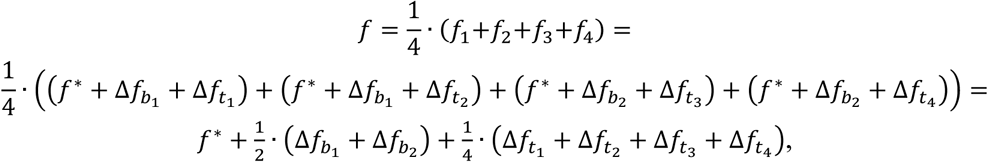

where *f*\* is the true frequency. Applying basic properties of variance the variance of mean frequency:

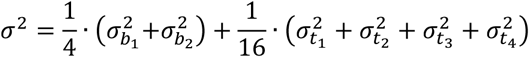

To estimate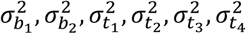 we used squared differences from pairwise comparison of replicas calculated aboves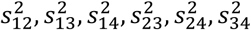:

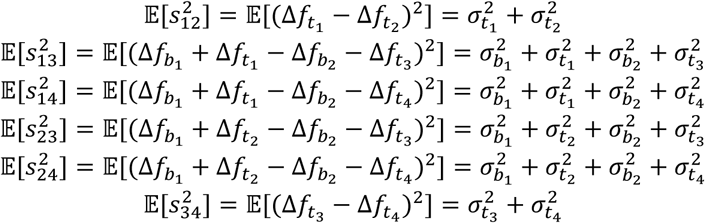

Therefore, the variance of mean frequency ***f*** can be found as:

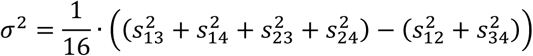

Recalling that variance and squared differences are a function of frequency:

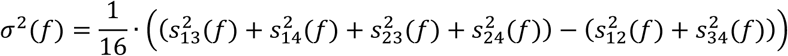

For each segment and time point we calculated the numerical function *σ*^2^(*f*). Then for each genotype having mean frequency *f*_*x*_ we estimated its variance as *σ*^2^ (*f*_*x*_)

### Merging amino acid genotypes

We merged nucleotide genotypes that corresponded to the same amino acid sequence and summed their frequencies and variances. We filtered out all genotypes *x* which had any of following patterns:

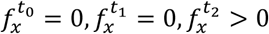 or 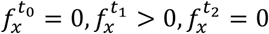 or 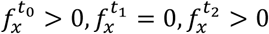.Fraction of such genotypes were <0.5% for all segments except S9, for which it was 4.5%

For further analysis, this amino acid dataset was used except when specified.

### Fitness estimation

Number of cells in a pool with particular genotype x after time interval ***t*** increases exponentially

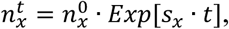

where ***s_x_*** is absolute fitness. Frequency of genotype x as well depends exponentially on absolute fitness with an additional multiplicative factor:

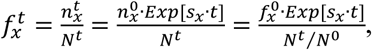

where *N*^*t*^ and *N*^0^ are total cell numbers in a pool at time points 0 and *t.* Factor 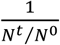 reflects the total growth of population, it changes with time but is the same for all genotypes. Therefore, we can rewrite genotype frequency at time *t* as:

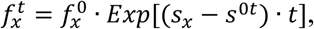

where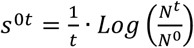

In the measured dataset for each genotype *x* we have 3 measurements of frequency 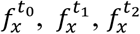 and their errors 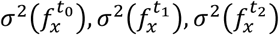. To estimate genotype fitness we minimized relative squared errors of exponential fit as function of fitness ***s_x_*** and initial frequency 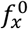:

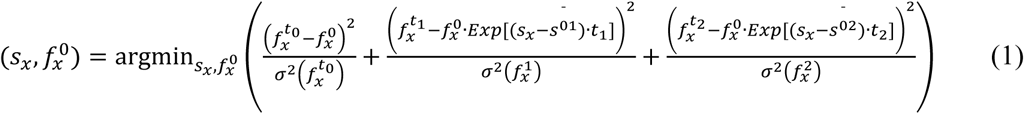

This formula contains four parameters common for all genotypes from one segment: *s*^01^*,s*^02^*,t*_1_*,t*_2_. Further we will perform additional shifting and scaling of fitness values (see next section), therefore, without loss of generality we could sets^01^ = 0 and *t*_1_ = 1.Ideally,t_2_/t_1_ should equal 14, however, we noticed that this ratio does not hold for many segments and fitted *k* = *t*_2_/*t*_1_ from data instead of using value 14.

To find specific *s*^02^ and *k* for each segment we selected genotypes with high frequencies at *t*_0_ (*t*_0_ >25 ⋅ 10^−6^) which corresponds to ~500-1000 reads per technical replicate. Each segment contains 10^3^-10^4^ genotypes that meet this criterion. We minimized eq. (1) for selected genotypes trying all possible combinations of (*s*^02^*,k*) from a grid where s^02^*ε*[0,1.2] with step 0.01 and *kε*[1,14] with step 0.1 and choose (*s^02^,k*) which gives minimal (*).

Finally, given (*s*^02^, *k*) for each segment we found *s*_*x*_ for each genotype. Errors for fitness values, 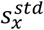, were estimated as standard error of best-fit parameter.

For genotypes with frequencies pattern 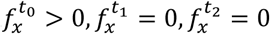 fit of eq. (1) cannot be obtained. Therefore we defined upper boundary for their fitness value as 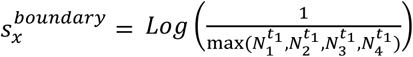, where 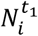, *i* 1..4 are total read numbers at time point *t*_*1*_ in *i* replica.

### Fitness rescaling

We scaled fitness such that lethal genotypes have fitness 0 and neutral genotypes have fitness 1. We assumed that genotypes with a stop codon or frame shift are lethal. Thus, for each segment we linearly rescaled the fitness distribution so that 95% of genotypes with nonsense mutations have a fitness of 0 and so that the local maximum of the fitness distribution of genotypes with extant amino acids is 1. The scaling around the local maximum led to the shift of fitness values of less than +/- 0.025 in each of the 12 segments compared to the measured wildtype strains and did not affect our results (for scale, we called a substitution non-neutral if its effect on fitness was > 0.4). All fitness values which became smaller than 0 were set to 0.

### Quality control and comparison of synonymous sequences

We used nucleotide synonymous sequences as an internal control. The error rate for a measurement of fitness of an amino acid sequence depends on the number of synonymous sequences, *n,* that were used to estimate it. Therefore, we estimated the false discovery rates separately for categories with *n=1,..10* variants. For each amino acid genotype with more than *n* synonymous variants we merged random combination of *n* of its nucleotide genotypes and estimated fitness. We then calculated the difference between this fitness and the fitness of the corresponding amino acid sequence. We classified case as “false unfit” if difference was <-0.4 and as “false fit” if difference was >0.4. The fraction of such cases gives us false discovery rates for genotypes having *n* synonymous variants. To get total false discovery rates for each segment we averaged “false unfit” and “false fit” rates for different *n* with weights equal to the fractions of genotypes in amino acid dataset which have *n* synonymous variant (**Supplementary Table 1**). The high correlation between biological replicas (**Supplementary Table 6**) confirms high accuracy of our high-throughput experiments, with the exception of segment 9.

## Predicting fitness using deep learning

To predict the unidimensional fitness function based on additive contribution of extant amino acid states we used deep learning, a powerful machine learning technique, capable of constructing virtually any function, even with a simple neural network architecture. To convert amino acid sequences into a binary feature matrix we used one-hot encoding strategy, in which each feature (column in the matrix) indicates the presence or absence of a particular amino acid state.

To optimise the accuracy/overfitting ratio, we tested over a hundred of different neural network architectures and parameters. As a starting point, we selected a number of complicated architectures, which describe our data, but are prone to overfitting due to a large number of parameters. We then gradually reduced the number of layers and neurons to reduce the overfitting, while controlling for accuracy.

Our final architecture consists of three layers and 22 neurons in total (**Supplementary Fig. 4**). Each of the neurons performs a linear transformation of the input and subsequently applies a non-linear activation function (a sigmoid) to the result. The output of the first layer, therefore, is a sigmoid of a linear transformation of the feature vector (c_1_^T^x). The second layer decompresses the hidden nonlinear representation into 20 sigmoids, the combination of which is further linearly transformed with the only neuron of the third layer and wrapped into another sigmoid function:

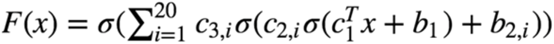

where 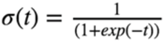 *X*- is the feature vector, *c*_1_ - the vector of coefficients, corresponding to the first layer, *c*_*n*_,_*k*_ - coefficient corresponding to the *n*-th layer and the *k*-th neuron, *b*_*n*_ - bias corresponding to the *n*-th layer. Crucially, this relatively simple architecture is capable of fitting virtually any function^53^, thus, in contrast to conventional logistic regression, in our approach we select the correct model from a vast variety of functions.

The key idea of our approach is that the number of neurons in the first layer of the neural network determines the number of linear combinations of mutations (or fitness potentials) used in order to predict mutant fitness. In other words, each neuron in the first layer assigns a single unique weight to every amino acid state in the dataset (**Supplementary Fig. 4**). Multiplication of such weight vectors and binary genotype vectors result in fitness potential. Thus, the number of neurons in the first layer of the architecture basically determines the dimensionality of epistasis we assume. The obtained fitness potentials are then transformed by a nonlinear phase shift function constructed by the 22 neurons of the neural network.

The architecture simplicity avoids overfitting, which was further prevented by using early stopping and keeping 10% of data as a test set. The loss function that is being optimised in our experiments is not convex, which leads to a high probability of getting stuck in different local minima. To ensure reproducibility, each of our models was constructed ten independent times using random train-test splits.

Each model was trained for under 100 epochs using mean squared error as the loss function. An unpublished adaptive learning rate method proposed by Geoff Hinton, RMSProp, was used as the optimiser. This algorithm is a version of a mini-batch stochastic gradient descent, utilising the gradient magnitude of the recent gradients in order to normalise the current ones. All the weights were initialised using Xavier normal initialiser^54^.

## Paths between pairs of fit genotypes

For analysis in **Fig. 6d**, we first choose two fit “parental” genotypes, one randomly chosen genotype (eg: ABE) and the other parental genotype that is either *S. cerevisiae* wildtype genotype (inter-segmental) or another random fit genotype in the data (intra-segmental) (eg: abe). The two genotypes in this example are Hamming Distance 3 apart (HD=3). We next compute all (HD2-2) intermediate genotypes (eg: AbC, aBc, *et cetera)* and retain the subset that were experimentally measured. We represent the two parental genotypes and all measured intermediate genotypes as an undirected graph in which each genotype is a vertex. All genotypes one substitution apart are connected by an unweighted edge. The shortest possible path for a given pair of genotypes is of length HD. We find all shortest paths between the two parental genotypes using a breadth-first search. We next remove all vertices (genotypes) that are unfit, and recompute the number of shortest between the two parental genotypes. For example, in **Fig. 6a**, there are six paths of length three if you take into account all genotypes, but only three paths of length three if you take into account only fit genotypes.

## Clustering of unfit genotypes in sequence space

For the analysis in **Fig. 6e**, we first represent the two parental genotypes and all measured intermediate genotypes as an undirected graph in which each genotype is a vertex. All genotypes one substitution apart are connected by an unweighted edge. We can then compute the degree (number of genotypes of distance one) for each vertex (genotype). We do so randomly drawing from all measured genotypes and using only unfit genotypes or using the same number but randomly chosen genotypes. For the randomly chosen genotypes, the value is the average over 1000 runs.

## Quantifying sign epistasis

For each substitution (eg: C -> S at position 141), we considered only those that exhibit a large fitness effect (abs. difference > 0.4) comprising a set of substitutions with large effects. For each substitution we divided the genetic backgrounds into two categories: those in which the substitution caused a > 0.4 increase in fitness, and those backgrounds in which the substitution caused > 0.4 decrease in fitness. A single substitution can cause a large increase in fitness in some backgrounds and a large decrease in others due to two possible reasons: sign epistasis or experimental error. To differentiate the two cases, we identified secondary substitutions that significantly alter the ratio of large increases to large decreases in fitness (Fisher’s exact test, Bonferroni corrected p-value < 0.05). We only consider a site to be under sign epistasis if there is a second site that alters the frequency of sign epistasis in a statistically significant manner, i.e. more frequently than expected by chance alone.

## Structural analysis

### Structure prediction

An initial model was obtained with the I-TASSER server^55^. The list of top 10 PDB structural templates picked up by the I-TASSER included high-quality crystal structures of imidazoleglycerol-phosphate dehydratases from *Arabidopsis thaliana* and *Cryptococcus neoformans.* Coordinates of the top-scoring model (C-score=0.21, estimated TM-score = 0.74±0.11, estimated RMSD = 5.1±3.3Å) and the predicted normalized B-factor^56^ were used for further analysis. The value of the model quality metric (TM-score >0.5) indicates a model of correct topology. The proteins structurally close to the final model (RMSD 0.6 - 1.7Å are PDB IDs 4MU0, 4GQU, 1RHY, 5DNL and 2AE8 from *Arabidopsis thaliana, Mycobacterium tuberculosis, Cryptococcus neoformans, Pyrococcus furiosus* and *Staphylococcus aureus.*

We measured the distribution of distances (in angstroms) between pairs of residues that exhibit strong sign epistasis (**Supplementary Table 1**, ReallyPositivePair == TRUE), and compared it with the distribution of pairwise distances among residues for which we have sufficient data to be certain that a given pair does not exhibit sign epistasis (**Supplementary Table 1**, ReallyNegativePair == TRUE).

### ΔΔG prediction

Cartesian_ddg application^57^ from Rosetta version 2017.08.59291 was used for ΔΔG predictions. Top-scoring I-TASSER model was pre-minimized using the Relax^58^ application in dual-space^59^ with the flags: -relax:dualspace true; -ex1; -ex2; -use_input_sc; -flip_HNQ; - no_optH false; -relax:min_typelbfgs_armijo_nonmonotone; -nonideal. The best scoring model from 1000 structures was selected. The effect of up to 4 mutations (54,500 genotypes in total) was assessed in Cartesian space with the Talaris_2014 score function, and the -fa_max_dis 9.0 flag. ΔΔG was estimated as a difference of mean score for 3 independent runs for every mutant and the wild-type score.

**Supplementary Figure 1.**
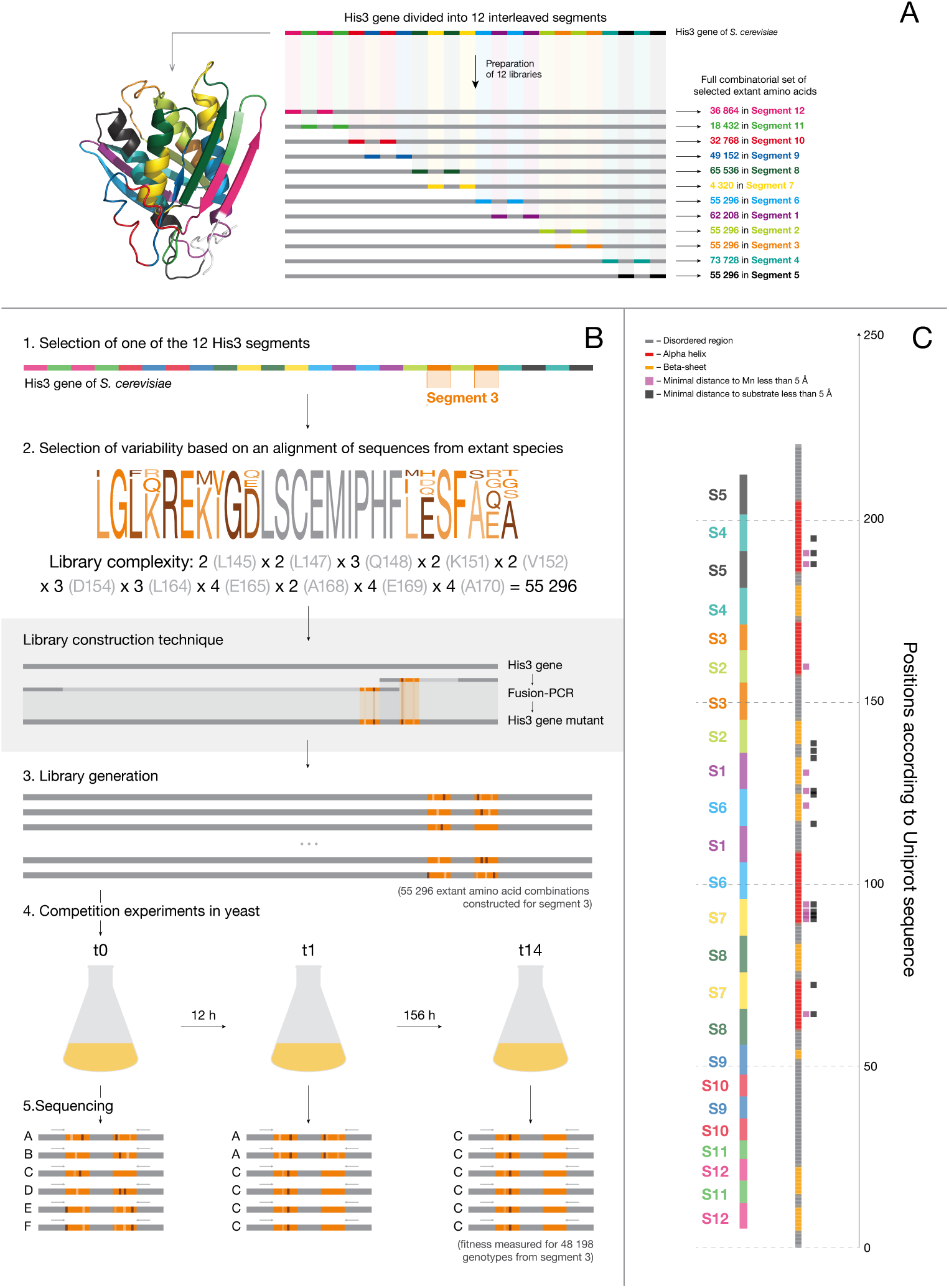
Experimental design. **a,** The sequence of the His3 protein from S. cerevisiae was separated into 12 independent segments of similar lengths, such that the full combinatorial set of extant amino acid substitutions was less than 100,000 possible genotypes. These segments represented different combinations of structural elements of the His3 protein structure. **b,** For each of the 12 segments from His3, we selected extant amino acid states using a multiple alignment of His3 orthologues from 396 species, preferentially incorporating states from 21 yeast species, the variability is shown in segment 3 as an example. Mutant degenerate codon libraries were constructed by fusion PCR of two synthesized variable halves of each segment. These high-complexity plasmid libraries were transformed into haploid His3 knockout S. cerevisiae strain. The growth rate of yeast carrying different extant amino acid state combinations in His3 gene was measured using serial batch culture in the absence of histidine with 12 hours between ~100-fold dilutions. To estimate the fitness of yeast mutants their relative abundance was measured at three points: in the initial population before selection (t0), in the population after 12 hours of growth in the selective medium (t1), and in the final population after 168 hours of growth in the selective medium (t14). To assess the fitness of individual mutants the segments from three populations were amplified and sequenced. The relative abundance of each sequence was used as a proxy for abundance of the associated yeast mutant, which in turn determines its fitness. **c**, Secondary structure of His3 mapped to the segments in our experiments.

**Supplementary Figure 2.**
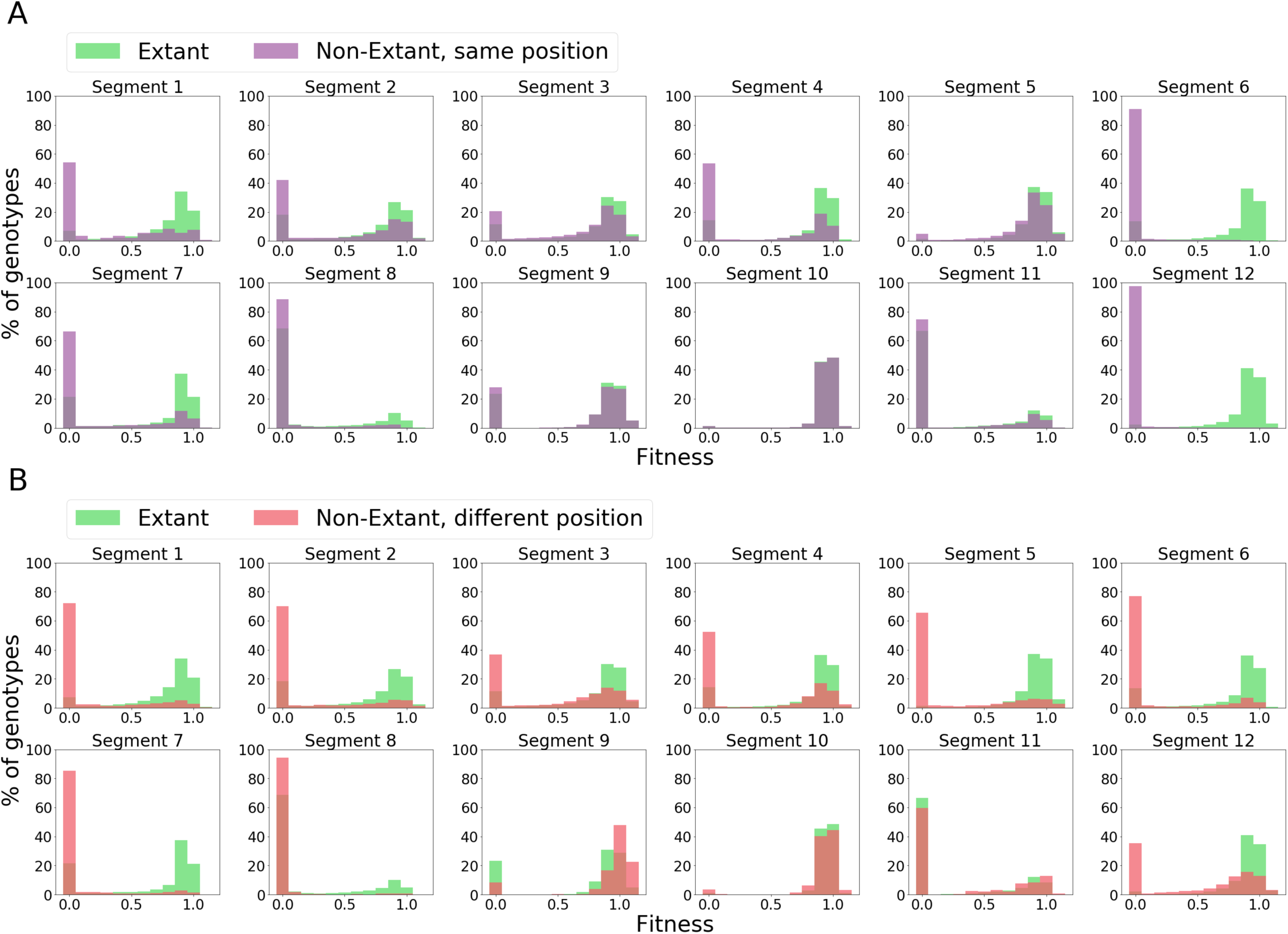
Segment-specific fitness distributions for extant and non-extant amino acid states. **a,** The fitness distribution for each segment for genotypes consisting only of extant amino acid states (green) or that contain one or more non-extant amino acid states (purple) only at positions with a substitution in the extant library. **b,** The fitness distribution for each segment for genotypes consisting only of extant amino acid states (green) and genotypes with mutations at other positions in that segment (red).

**Supplementary Figure 3.**
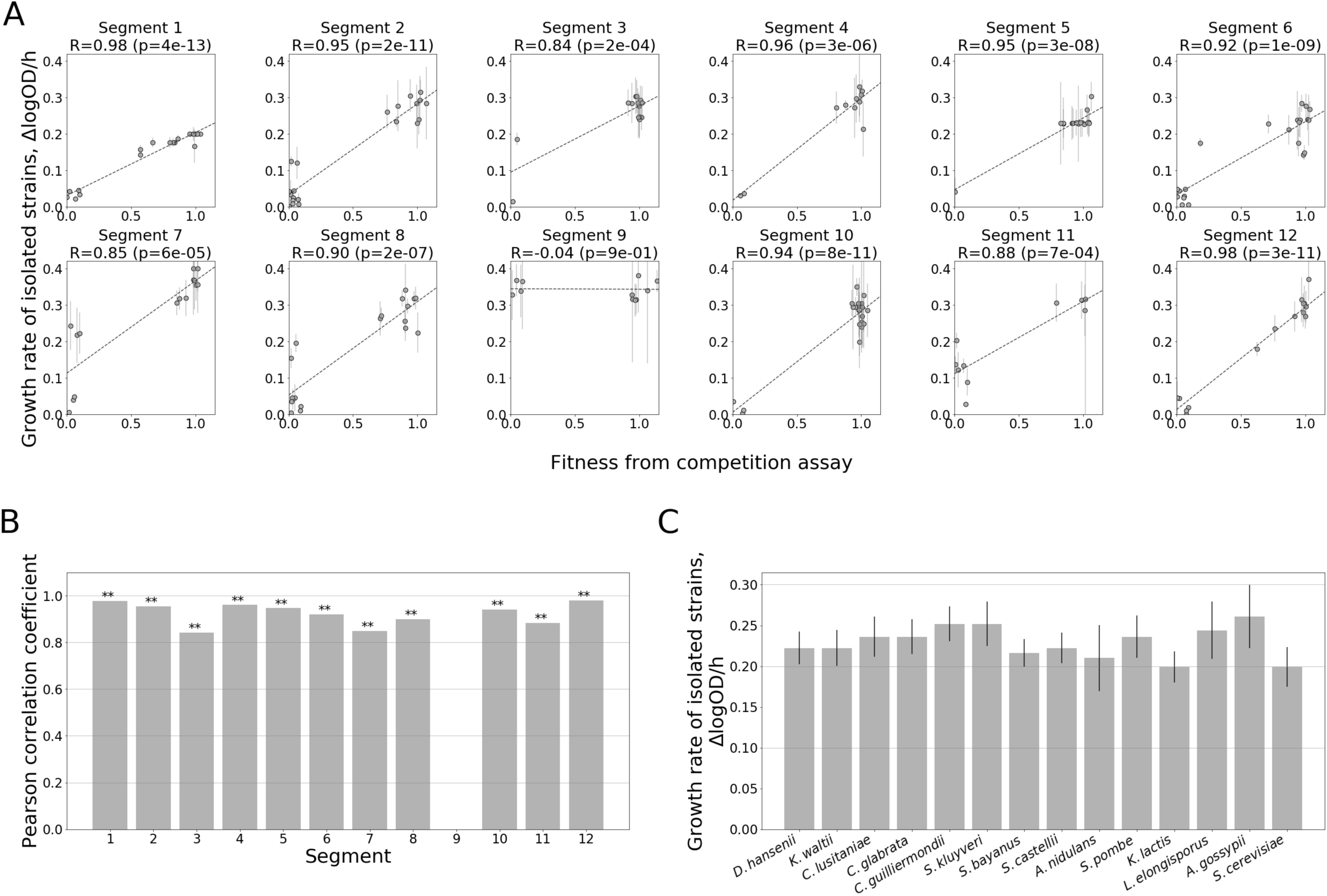
Growth rate measurement of isolated strains. **a,** Comparison of fitness values from the pooled competition assay with growth rates of isolated strains as measured in a microplate reader. Error bars for growth rates show s.e.m. of 6 replicates. **b**, Pearson correlation coefficients between fitness values from competition and growth rates of isolated strains for each segment. ** signifies p-value < 0.005 (correlation test). **c**, His3p orthologues from different species complement a Δ his3 deletion in *S. cerevisiae.* Growth rates of transformants containing whole HIS3 orthologous genes from other yeast species. Error bars for growth rates show s.e.m. of ≥ 7 replicates.

**Supplementary Figure 4.**
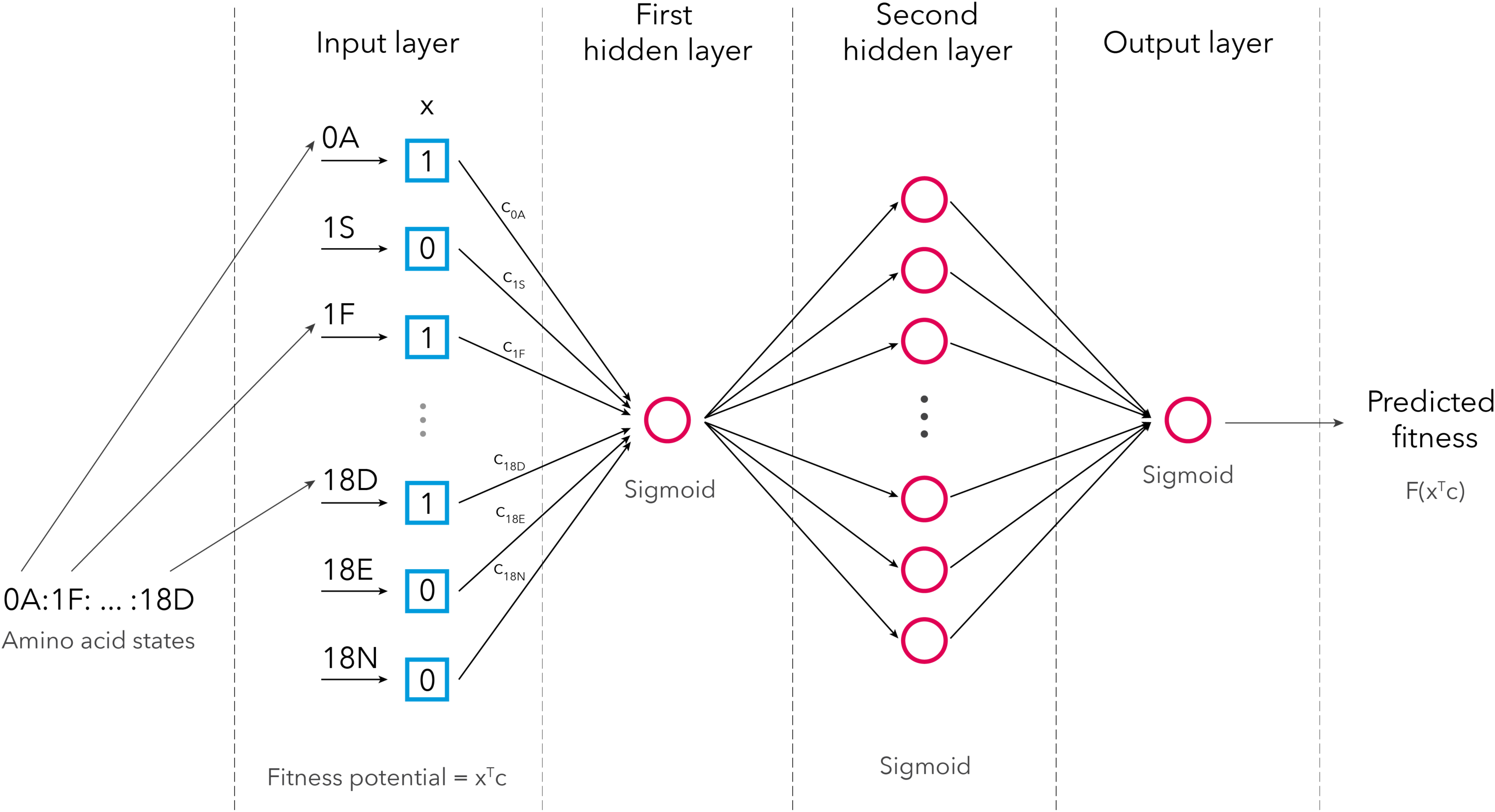
Schematic representation of the deep learning approach. Each genotype was encoded as a binary vector (x). During training, each of the substitutions was assigned a coefficient (*c*_*i*_), comprising a vector of coefficients (c). The multiplication of these two vectors is the fitness potential of the genotype. After going through three layers, each with a sigmoid activation function, the predicted fitness is obtained.

**Supplementary Figure 5.**
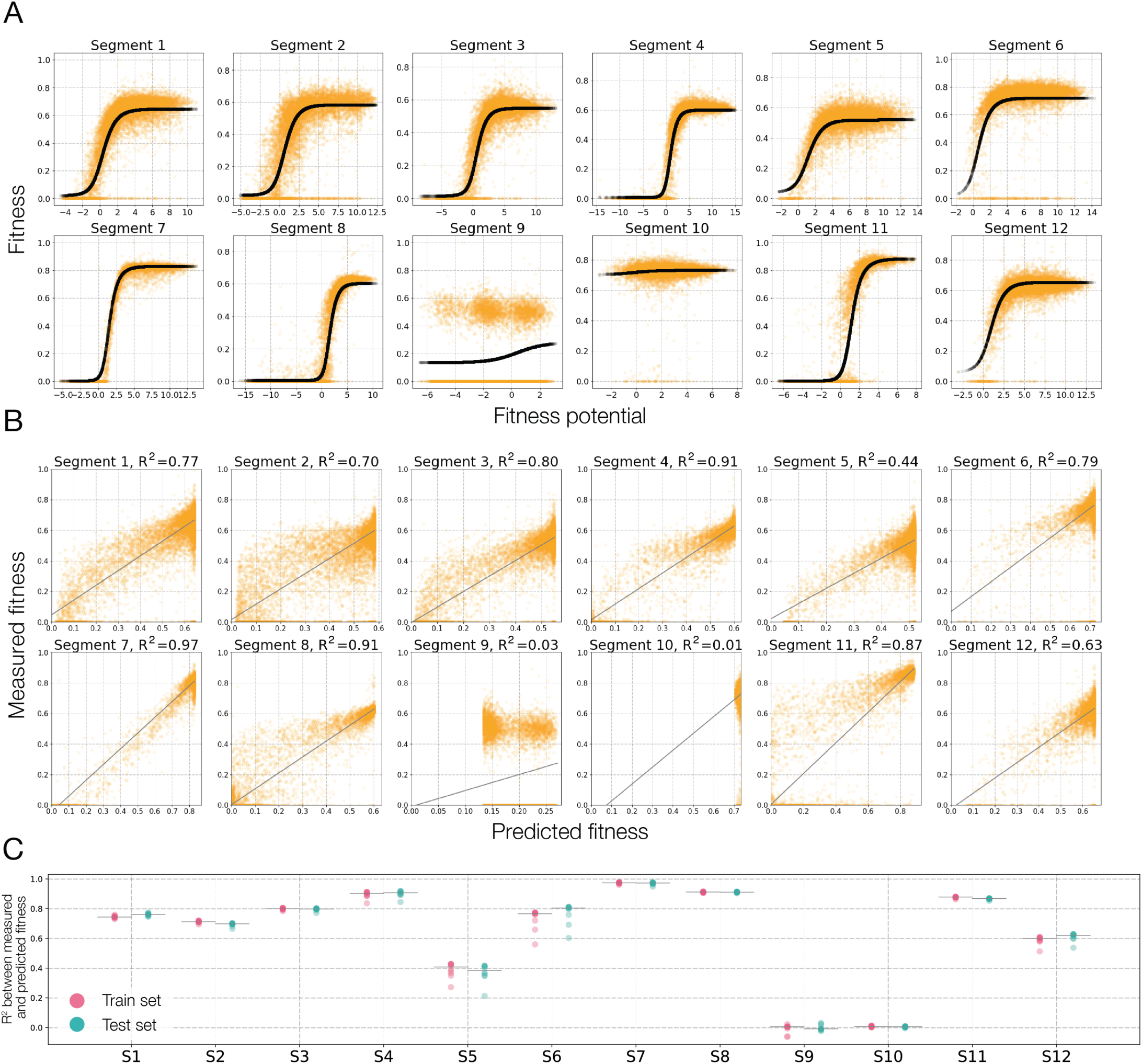
The fitness potential of the His3 fitness landscape. **a**, Fitness potential predicted by the neural network as a function of the measured fitness for all 12 segments. **b**, The correlation between the fitness predicted by the fitness potential and the measured fitness. **c,** Training and test R^2^ for each segment for 20-fold cross-validation.

**Supplementary Figure 6.**
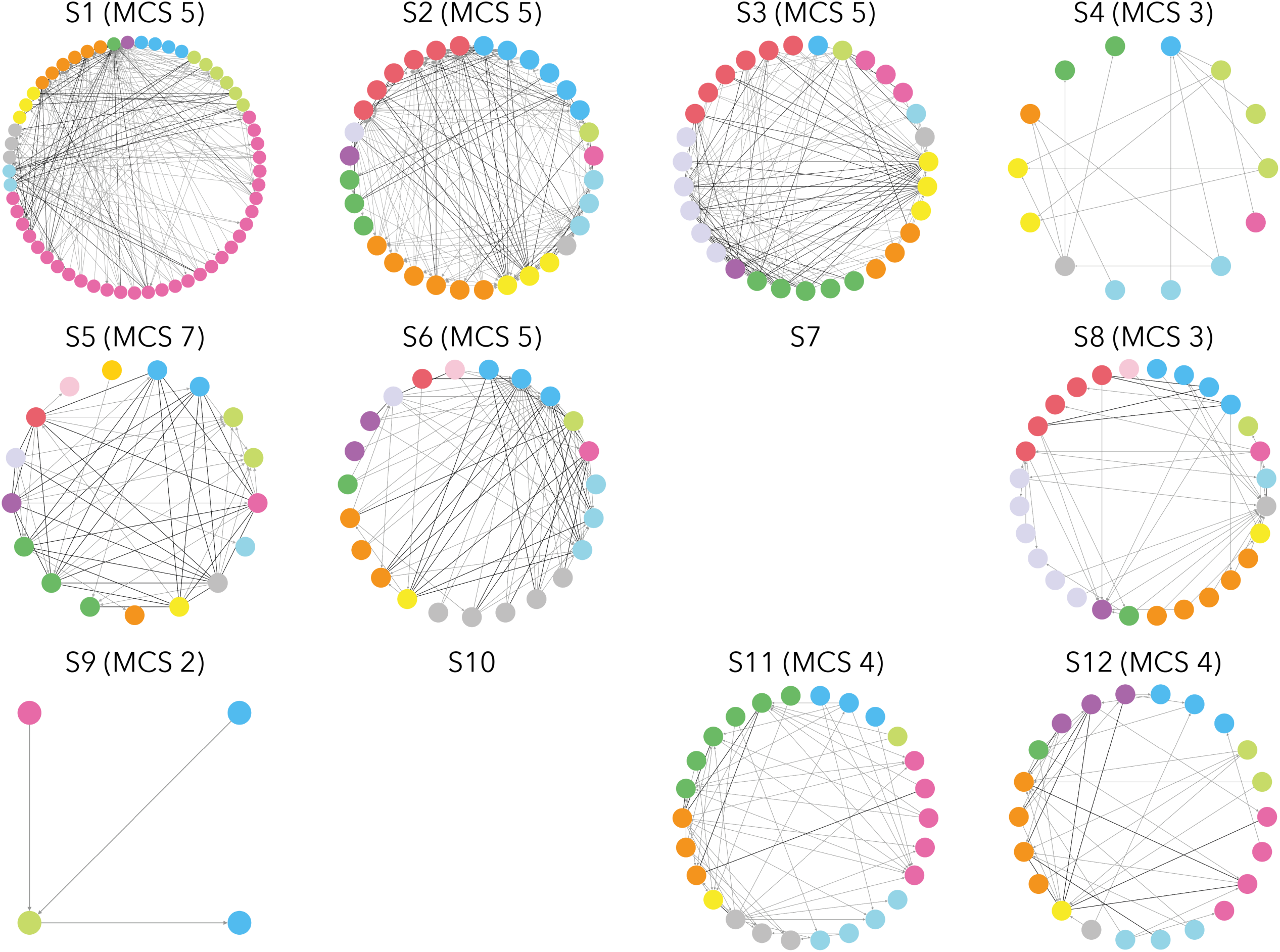
Sign epistasis dimensionality graphs for all twelve segments. Each node represents a substitution, with multiple substitutions at the same site having the same colour. Substitutions under reciprocal sign epistasis are indicated by black lines while grey arrows indicate unidirectional sign epistasis.

**Supplementary Figure 7.**
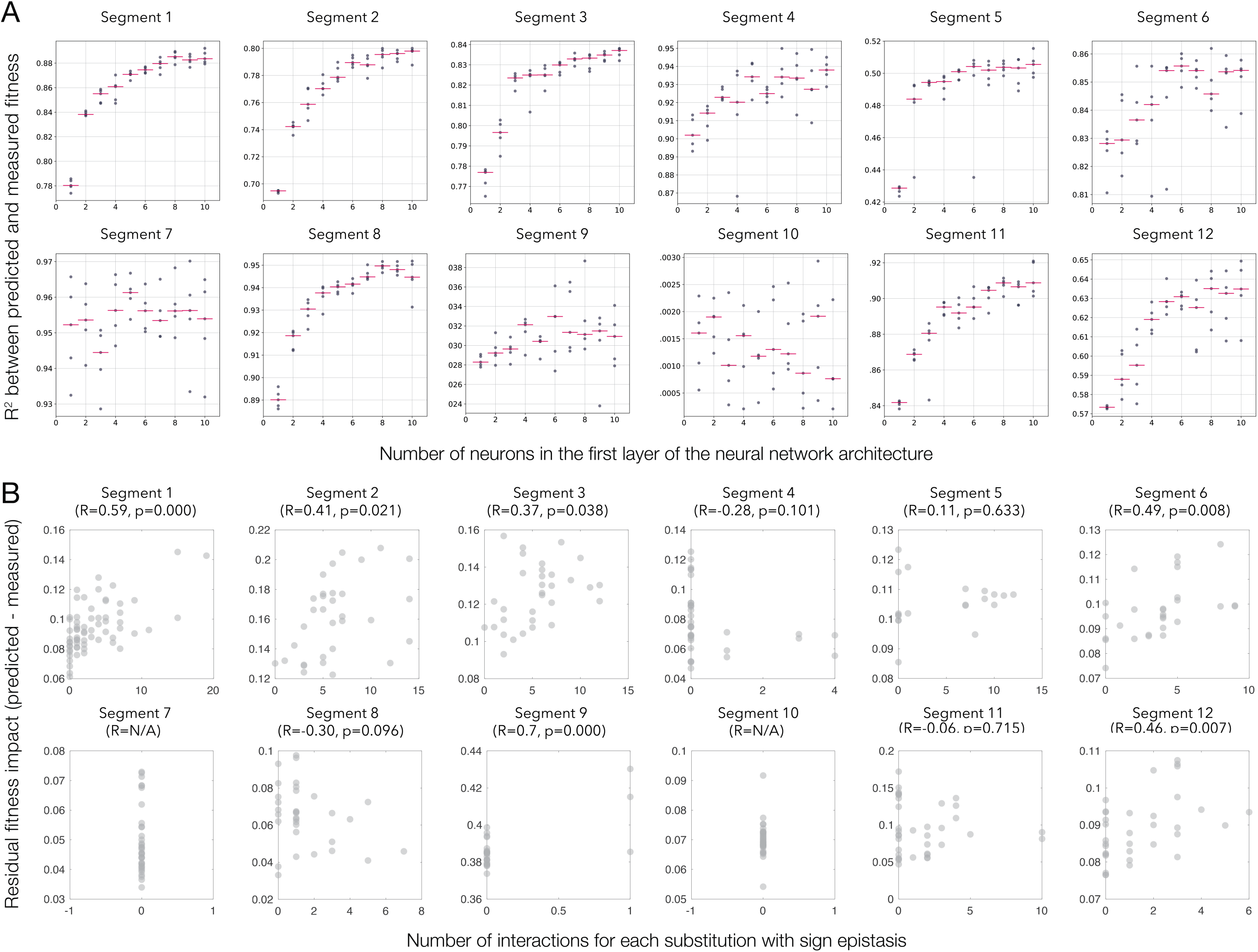
Multidimensional description of epistasis in His3 segments. **a,** Increasing the number of neurons in the first layer of the neural network, which is equivalent to increasing the number of underlying fitness potentials, leads to more accurate models for segments with detected sign epistasis. Each dot corresponds to an independent optimization of model parameters. **b,** Number of sign epistatic interactions of certain substitutions against average model prediction power for mutants including these substitutions.

**Supplementary Figure 8.**
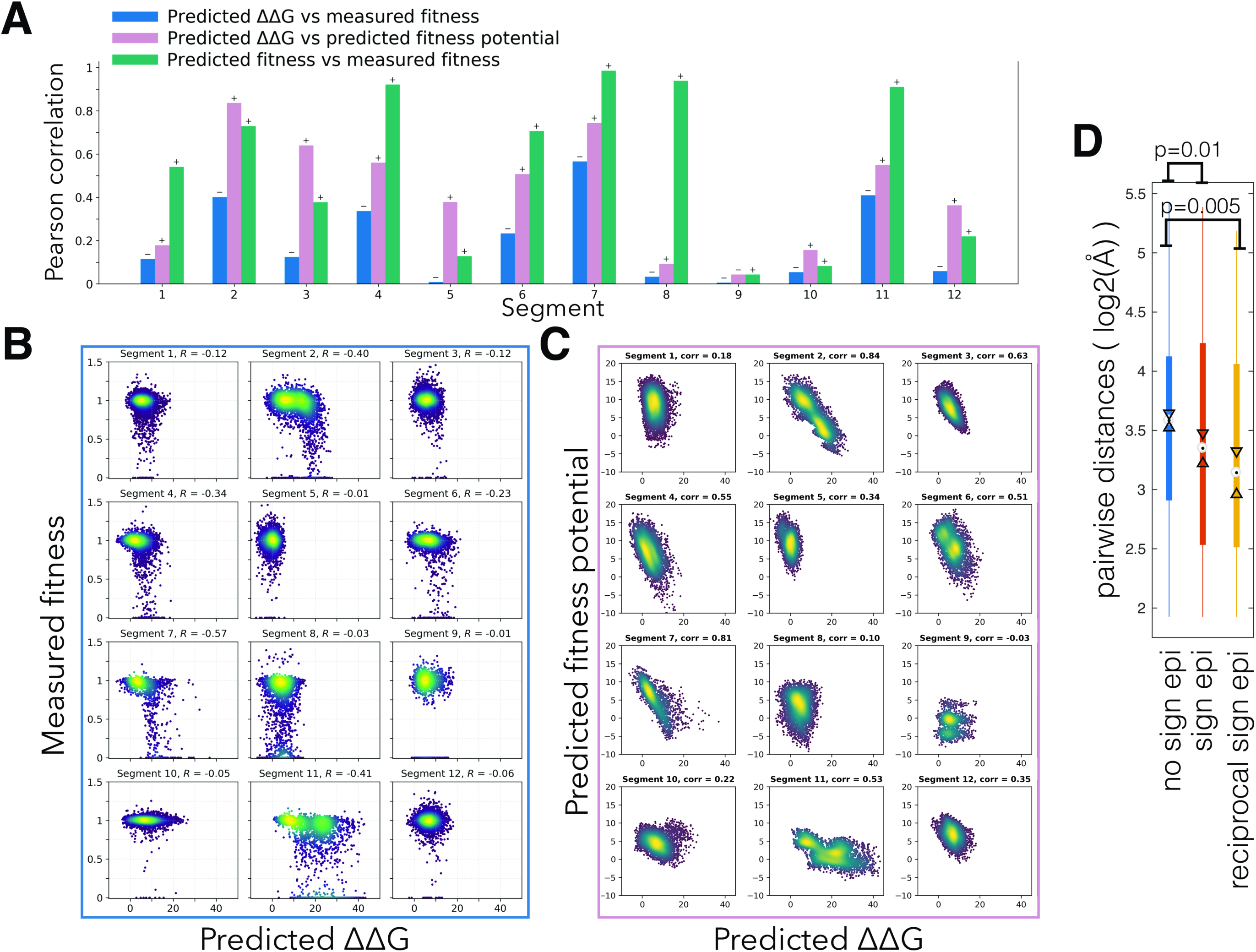
Protein stability and the fitness potential. **a,** A comparison of correlation coefficients between predicted and measured values across segments. **b,c**, correlations between the estimated impact of substitutions on free energy (ΔΔG), fitness potential and fitness. ΔΔG correlates better with fitness potential than with fitness. **d**, Pairs of sites that exhibit sign (connected by a light edge in Supplementary Figure 6) and those that exhibit reciprocal sign epistasis (connected by a dark edge in Supplementary Figure 6) are closer together in the His3p structure that randomly chosen non-connected pairs of positions that exhibit sign epistasis.

**Supplementary Figure 9.**
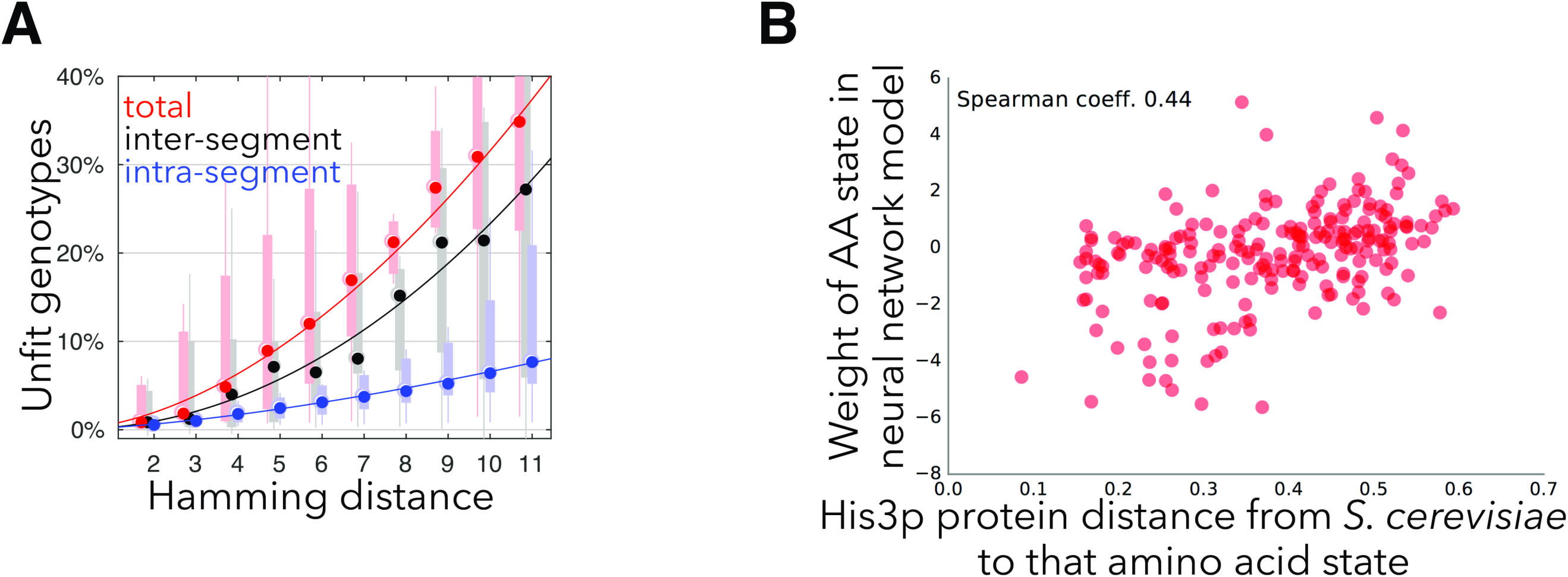
Decoupling inter- and intra-segmental epistasis. **a,** The fraction of unfit genotypes between *S. cerevisiae* and any other genotype consisting of extant amino acid states with high (blue) or any (red) fitness, and genotypes in the latter but not the former category (black) as a function of the Hamming distance between the two boundary genotypes. Points indicate median, the bars and lines indicate 50% of the genotypes and genotypes 2.7 sigmas from the mean, respectively. **b,** The neural network model assigns higher weights to amino acid states that that first occur in His3 orthologues farther from *S cerevisiae,* indicating the presence of intrasegmental interactions.

**Supplementary Information 1. Multiple alignment of His3 orthologues.**

**Supplementary Information 2. Multidimensional description of epistasis in His3 segments.** Fitness as a function of two fitness potentials (black dots, measured fitness is depicted in orange).

